# 14-3-3 promotes sarcolemmal expression of cardiac Ca_V_1.2 and nucleates isoproterenol-triggered channel super-clustering

**DOI:** 10.1101/2024.08.16.607987

**Authors:** Heather C. Spooner, Alexandre D. Costa, Adriana Hernández González, Husna Ibrahimkhail, Vladimir Yarov-Yarovoy, Mary Horne, Eamonn J. Dickson, Rose E. Dixon

## Abstract

The L-type Ca^2+^ channel (Ca_V_1.2) is essential for cardiac excitation-contraction coupling. To contribute to the inward Ca^2+^ flux that drives Ca^2+^-induced-Ca^2+^-release, Ca_V_1.2 channels must be expressed on the sarcolemma; thus the regulatory mechanisms that tune Ca_V_1.2 expression to meet contractile demand are an emerging area of research. A ubiquitously expressed protein called 14-3-3 has been proposed to affect Ca^2+^ channel trafficking in non-myocytes, however whether 14-3-3 has similar effects on Ca_V_1.2 in cardiomyocytes is unknown. 14-3-3 preferentially binds phospho-serine/threonine residues to affect many cellular processes and is known to regulate cardiac ion channels including Na_V_1.5 and hERG. Altered 14-3-3 expression and function have been implicated in cardiac pathologies including hypertrophy. Accordingly, we tested the hypothesis that 14-3-3 interacts with Ca_V_1.2 in a phosphorylation-dependent manner and regulates cardiac Ca_V_1.2 trafficking and recycling. Confocal imaging, proximity ligation assays, super-resolution imaging, and co-immunoprecipitation revealed a population of 14-3-3 colocalized and closely associated with Ca_V_1.2. The degree of 14-3-3/Ca_V_1.2 colocalization increased upon stimulation of *β*-adrenergic receptors with isoproterenol. Notably, only the 14-3-3-associated Ca_V_1.2 population displayed increased cluster size with isoproterenol, revealing a role for 14-3-3 as a nucleation factor that directs Ca_V_1.2 super-clustering. 14-3-3 overexpression increased basal Ca_V_1.2 cluster size and Ca^2+^ currents in ventricular myocytes, with maintained channel responsivity to isoproterenol. In contrast, isoproterenol-stimulated augmentation of sarcolemmal Ca_V_1.2 expression and currents in ventricular myocytes were abrogated by 14-3-3 inhibition. These data support a model where 14-3-3 interacts with Ca_V_1.2 in a phosphorylation-dependent manner to promote enhanced trafficking/recycling, clustering, and activity during *β*-adrenergic stimulation.

**Significance Statement:** The L-type Ca^2+^ channel, Ca_V_1.2, plays an essential role in excitation-contraction coupling in the heart and in part regulates the overall strength of contraction during basal and fight-or-flight *β*-adrenergic signaling conditions. Proteins that modulate the trafficking and/or activity of Ca_V_1.2 are interesting both from a physiological and pathological perspective, since alterations in Ca_V_1.2 can impact action potential duration and cause arrythmias. A small protein called 14-3-3 regulates other ion channels in the heart and other Ca^2+^ channels, but how it may interact with Ca_V_1.2 in the heart has never been studied. Examining factors that affect Ca_V_1.2 at rest and during *β*-adrenergic stimulation is crucial for our ability to understand and treat disease and aging conditions where these pathways are altered.

## Introduction

The Ca_V_1.2 subtype of voltage-gated L-type Ca^2+^ channels play critical roles in excitation-contraction coupling in smooth and cardiac muscle, excitation-secretion coupling in endocrine cells, and in excitation-transcription coupling in neurons. In cardiomyocytes, depolarization of the sarcolemma by an action potential activates a subset of Ca_V_1.2 channels, allowing Ca^2+^ influx that activates nearby type 2 ryanodine receptors (RyR2) on the junctional sarcoplasmic reticulum (jSR). These channels facilitate a graded release of Ca^2+^ from the SR in a process known as Ca^2+^-induced Ca^2+^ release (CICR) and the overall rise in intracellular Ca^2+^ concentration triggers contraction (1). In skeletal muscle, contraction magnitude can be tuned by recruiting more, or less motor units, but in the heart every cardiomyocyte is activated during each beat of the heart thus there are no more cells to be recruited. Instead, the magnitude of cardiac muscle contractility is tuned by the degree of Ca^2+^ influx. Because Ca^2+^ release from the SR is graded, an increase in the trigger Ca^2+^ influx through Ca_V_1.2 also increases the Ca^2+^ transient amplitude and thus contraction magnitude. Accordingly, regulatory pathways, channel auxiliary subunits, and co-factors that can modulate Ca_V_1.2 channel activity and/or sarcolemmal expression are physiologically relevant and thus of particular interest. Furthermore, since alterations in Ca_V_1.2 channel activity can impact action potential duration and promote cardiac arrhythmia (2), alterations in regulatory proteins that modulate channel activity are also interesting from a pathological perspective.

Cardiac Ca_V_1.2 channels are multimeric protein complexes consisting of a pore-forming Ca_V_α_1C_ subunit, an auxiliary Ca_V_β predominantly Ca_V_β_2b_ in cardiomyocytes (3), and a Ca_V_α_2_δ subunit. Depending on the regulatory state of the channel the cardiac Ca_V_1.2 complex may also contain the RGK (<u>R</u>ad, <u>R</u>em, <u>R</u>em2, <u>G</u>em/<u>K</u>ir) protein Rad. These auxiliary proteins associate with the channel complex and have all been found to influence channel expression at the plasma membrane/sarcolemma (4–7). In addition, Rad has been shown to play an essential role in the functional upregulation of Ca_V_1.2 during *β*-adrenergic receptor stimulation when PKA-mediated phosphorylation of Rad disrupts its inhibitory interaction with the Ca_V_β auxiliary subunit resulting in increased channel activity (8–12). These findings underscore the importance of identifying and investigating the functional role of Ca_V_1.2 channel cofactors. In this study we present evidence supporting 14-3-3 as a novel cardiac Ca_V_1.2 channel interacting protein that affects responsivity to *β*-adrenergic receptor signaling.

14-3-3 is a small (∼30 kDa) ubiquitously expressed protein that has many diverse roles in subcellular processes including regulation of transcription, ubiquitination, and chaperoning (13). The numerical name of this protein originated when it was first identified in bovine brain lysates within the <u>14</u>^th^ elution fraction on a column following DEAE-cellulose chromatography, and for their subsequent electrophoretic mobility on a gel where they migrated to the <u>3.3</u> position. There are seven mammalian isoforms of 14-3-3 (β, γ, ε, ζ, η, θ, and σ), each encoded by separate genes *(YWHAB, YWHAG, YWHAE, YWHAZ, YWHAH, YWHAQ*, and *SFN* or *Stratifin*), with 14-3-3ε emerging as the most abundant cardiac isoform at both the protein (14) and mRNA level (15). 14-3-3 isoforms preferentially bind phospho-serine/threonines (16, 17) and are known to modulate the activity, trafficking/co-trafficking, or phosphorylation state of other cardiac ion channels including HERG (18), TASK-1 and TASK-3 (19), Na_V_1.5 (20, 21), Kir2.1 (22), K_ATP_ (15), SERCA (23), PMCA, (24, 25), and NCX (26). In addition, the trafficking and inactivation kinetics of the neuronal Ca_V_2.2 voltage-gated calcium channel is reportedly altered by 14-3-3 (27, 28). Despite the abundance of evidence for the role of 14-3-3 in regulation of other cardiac ion channels and regulators of calcium homeostasis, there is a scarcity of information on the effect of 14-3-3 on Ca_V_1.2 channels, although it is known that membrane-anchored 14-3-3ε (via fusion to a palmitoylated peptide) or membrane-recruitable 14-3-3ε (via fusion to a PKC C1 domain that translocates to the membrane upon PKC activation) affect Ca_V_1.2 activity implying that 14-3-3ε can interact with Ca_V_1.2 (29). Furthermore, 14-3-3 is known to interact with RGK proteins including Rad (30, 31), making it an interesting candidate for phosphorylation-dependent regulation of Ca_V_1.2 trafficking and activity.

Here we present an examination of the effects of 14-3-3 on multiple layers of Ca_V_1.2 trafficking and regulation. We report that 14-3-3 interacts with transiently transfected Ca_V_1.2 in tsA-201 cells and with endogenous Ca_V_1.2 channels within mouse ventricular myocytes where phosphorylation-dependent 14-3-3 interactions alter Ca_V_1.2 localization and nanoscale redistribution during *β*-adrenergic receptor stimulation. We report a role for 14-3-3 in regulating isoproterenol-stimulated Ca_V_1.2 channel activity and recycling finding that 14-3-3ε overexpression increases *I*_Ca_ amplitude and plasma membrane expression of Ca_V_1.2 in both tsA-201 cells and ventricular myocytes while treatment with a competitive inhibitor of 14-3-3 attenuates *β*-adrenergic receptor stimulated increases in *I*_Ca_ and abrogates isoproterenol stimulated channel recycling and superclustering. Based on our findings, we propose a model wherein 14-3-3 acts as a nucleation factor for Ca_V_1.2 clustering on the sarcolemma.

## Results

### 14-3-3 associates with Ca_V_1.2 in tsA-201 cells

To determine whether 14-3-3 associates with Ca_V_1.2, we performed co-immunoprecipitation (co-IP) assays to probe for Ca_V_1.2-14-3-3 interactions. We chose to focus on 14-3-3ε based on RT-PCR experiments showing 14-3-3ε has the highest mRNA expression in mouse and rat ventricular myocytes (15), and a study that found a membrane-targeted version of 14-3-3ε was able to modify Ca_V_1.2 activity in HEK-293 cells or cultured murine dorsal root ganglion (DRG) neurons, although the function and nature of this potential interaction and whether it applied to native 14-3-3 protein was not tested (29). We initially adopted a reductionist system and performed IPs on whole cell lysates from tsA-201 cells transiently transfected with FLAG-tagged Ca_V_α_1C_ and HA-tagged 14-3-3ε. The association of 14-3-3 with Ca_V_1.2 was assessed by immunoprecipitation (IP) of the tagged-proteins from the lysate followed by SDS-PAGE fractionation of the complexes and immunoblot (IB) analysis of the fractionated proteins. To assess the association of Ca_V_1.2 with 14-3-3ε, anti-HA (14-3-3ε) immunocomplexes were probed with an anti-FLAG (Ca_V_α_1C_) antibody (Fig. 1A; *top*). To determine whether the reverse, i.e. pull-down of Ca_V_1.2 would also bring down 14-3-3ε associated with it, the reciprocal IP using anti-FLAG followed by IB with anti-HA was performed (Fig. 1Α; *bottom*). In both cases, appreciable amounts of the putative complex partner were observed, confirming association of Ca_V_1.2 with 14-3-3ε in these cells.

**Figure 1.**
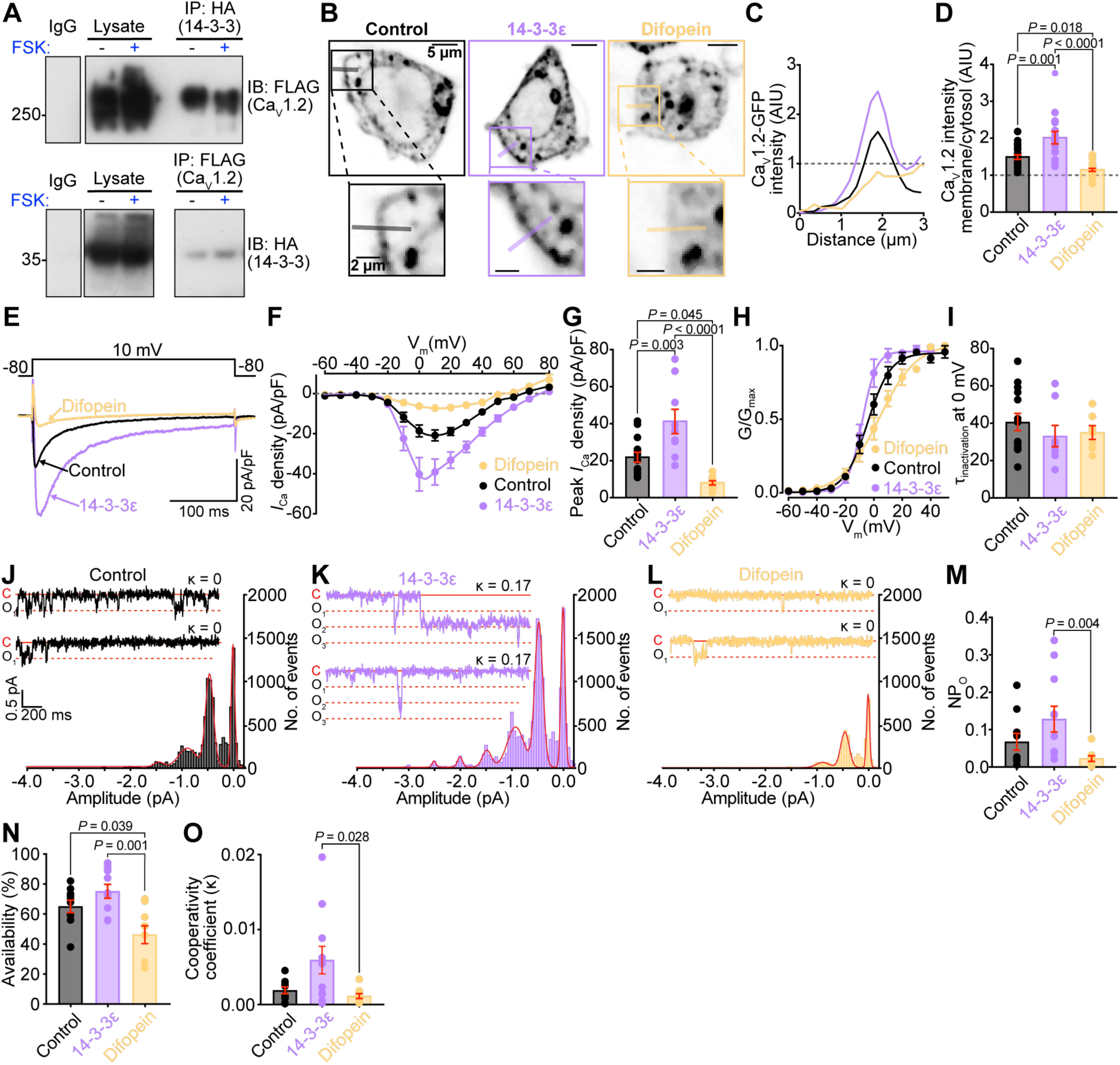
14-3-3 and Ca_V_1.2 co-immunoprecipitate and 14-3-3 levels regulate Ca_V_1.2 surface expression and whole cell calcium currents in tsA-201 cells. **A**, Immunoblots showing IgG control, lysate, and 14-3-3 IP lanes probed for Ca_V_1.2 and Ca_V_1.2 IP lanes probed for 14-3-3 in tsA-201 lysates treated with FSK (+), or with vehicle alone (−). B, representative confocal images showing Ca_V_1.2-RFP localization in control, 14-3-3ε overexpressing, or difopein-expressing tsA-201 cells. C, Plot profiles of membrane/cytosolic RFP intensity across the line regions-of-interest indicated in B. D, histogram summarizing the ratio between membrane and cytosolic Ca_V_1.2-RFP intensity (control: *n* = 22; 14-3-3ε: *n* = 15; difopein: *n* = 20). E, representative whole-cell currents elicited from transfected tsA-201 cells from each condition. F, I-V plots and G, histogram summarizing peak *I_Ca_* density (control: *n* = 14; 14-3-3ε: *n* = 9; difopein: *n* = 9). H, voltage-dependence of the normalized conductance (G/Gmax) fit with Boltzmann functions (control: *n* = 13; 14-3-3ε: *n* = 8; difopein: *n* = 7). I, histogram showing time constant (τ) of inactivation at 0 mV. J-L, All-points histograms with inset representative *i_Ca_* traces with corresponding cooperativity coefficient (κ) for control (J, *n* = 10, 474 sweeps), 14-3-3ε (K, *n* = 11, 481 sweeps), and difopein cells (L, *n* = 9, 448 sweeps). Amplitude histograms were fit with multi-component Gaussian functions (red lines). M-O, Summary data of NP_O_ (M), channel availability (N), and average κ per cell (O) (control: *n* = 10; 14-3-3ε: *n* = 11; difopein: *n* = 9). Data were analyzed using one-way ANOVAs (D, G, I, N, and O), or Kruskal-Wallis tests (M) with multiple comparison post-hoc tests. Error bars indicate SEM.

14-3-3 preferentially binds to phosphorylated serine or threonine residues on various proteins via electrostatic interactions (16, 32–34). In HERG K^+^ channels, another cardiac ion channel, phosphorylation-dependent interactions with 14-3-3 have been reported to shield phosphorylated residues from phosphatases thus prolonging *β*-adrenergic receptor (*β*-AR) stimulation of HERG activity (18). Recently a similar mechanism also has been shown for phospholamban, the small regulatory protein that confers PKA regulation on the cardiac SR calcium pump SERCA (23). To determine whether Ca_V_1.2-14-3-3 interactions can occur in a phosphorylation-dependent manner, co-IPs were performed on tsA-201 cell lysates, transfected as described above and treated with the adenylyl cyclase activator forskolin (FSK; 10 µM for 8 mins). No obvious FSK-stimulated increase in Ca_V_1.2-14-3-3 interactions was detected (Fig. 1A), however we cannot rule out that HA-14-3-3 is already associated with phosphorylated residues on Ca_V_1.2 in the control (no FSK) lysates and that our assay is not sensitive enough to detect additional association with FSK, or that direct 14-3-3-Ca_V_1.2 interactions are dependent on phosphorylation by a kinase not stimulated by FSK. It is also important to note that while tsA-201 cells do not endogenously express Ca_V_1.2, they do express significant amounts of 14-3-3 (35). Thus, the interactions observed here are likely an underestimation since they detect only overexpressed HA-tagged 14-3-3 in complex with Ca_V_1.2, and are blind to endogenous 14-3-3-Ca_V_1.2 complexes.

### Surface expression of Ca_V_1.2 is promoted by 14-3-3

Having established that 14-3-3 and Ca_V_1.2 associate with one another, we next investigated whether these interactions could promote surface trafficking of Ca_V_1.2 channels as has been reported for N-type Ca_V_2.2 channels (28). To test that idea, we used tsA-201 cells transiently transfected with the channel (Ca_V_α_1C_-RFP) and auxiliary subunits (Ca_V_β_2b_ and Ca_V_α_2_δ). In some experiments cells were additionally transfected with 14-3-3ε to achieve overexpression, or with a plasmid (pSCM138) encoding a YFP-tagged 14-3-3 inhibitor called difopein (for <u>di</u>meric <u>fo</u>urteen-three-three <u>pe</u>ptide <u>in</u>hibitor; a doublet of R18 inhibitory peptide; pSCM138) to compete for 14-3-3 binding or as a control, the plasmid pSCM174 encoding an inactivated form of the difopein peptide (a YFP-fused doublet of R18 containing D12K and E14K mutations) (36). After 48 hours, we measured surface localization of RFP-tagged Ca_V_1.2 using confocal microscopy (Fig. 1B-D). The localization of Ca_V_1.2 at or close to the plasma membrane was significantly reduced in cells co-transfected with difopein when compared to no difopein or inactivated difopein control cells (Fig. 1B-D, *SI appendix* Fig. S1A-B). Channels appeared to be stuck in intracellular compartments in difopein-expressing cells implying impaired trafficking processes. We found no significant difference in Ca_V_1.2 surface localization in the no difopein controls as compared to those expressing inactive difopein (*SI appendix* Fig. S1A-B) and thus we pooled those cells in all subsequent analyses. To determine whether this role in Ca_V_1.2 trafficking facilitation extended beyond 14-3-3ε and applied to other 14-3-3 isoforms, we repeated this experiment with multiple 14-3-3 isoforms (*SI appendix* Fig. S1B-C). While trending increases in Ca_V_1.2 plasma membrane localization were observed with every 14-3-3 isoform examined, the only statistically significant change was with 14-3-3ε, providing additional confidence in our decision to focus on 14-3-3ε.

If 14-3-3 does promote surface expression of Ca_V_1.2, then two logical predictions are: i) 14-3-3 overexpression should increase whole cell calcium current (*I_Ca_*), and ii) disruption of 14-3-3 -Ca_V_1.2 interactions with difopein should decrease the number of functional channels at the surface thus reducing *I_Ca_*. To test those predictions, we measured *I_Ca_* in tsA-201 cells transfected as described above and saw a 1.88 ± 0.30-fold increase in *I_Ca_* with 14-3-3ε overexpression, and very little current in cells with difopein (Fig. 1E-G). We did not observe significant changes in the voltage at half maximal activation (V_1/2_), a measure of voltage-dependence of activation or in inactivation kinetics with any perturbation (Fig. 1H-I, *SI appendix* Table S1). Since *I*_Ca_ is given by the product of the number of active channels, their open probability, and their unitary current (*i*_Ca_) amplitude (*I_Ca_* = *N* × *P_o_* × *i_Ca_*) we sought to elucidate which of these parameters were altered to explain the changes in *I*_Ca_ with 14-3-3ε overexpression or competitive inhibition. Cell-attached patch single channel recordings were thus performed with currents evoked using step depolarizations to −30 mV with Ca^2+^ as the charge carrier. These experiments revealed that control (or inactivated difopein expressing), 14-3-3ε overexpressing, and difopein expressing cells all had similar unitary current (*i*_Ca_) amplitudes (control −0.482 ± 0.018 pA, 14-3-3ε −0.500 ± 0.014 pA, difopein −0.444 ± 0.013 pA; Fig. 1J-L). Aligning with our *I*_Ca_ results, a trending increase in single channel activity (NP_o_) occurred with 14-3-3ε overexpression, while difopein significantly reduced NP_o_ (Fig. 1M). Thus, in the absence of changes in the unitary conductance, the effects of 14-3-3 on *I*_Ca_ appear to occur due to alterations in single channel activity.

### 14-3-3 augments cooperative interactions of Ca_V_1.2 channels

All-points histograms compiled from multiple cells (control *n* = 10 cells, 474 sweeps; 14-3-3ε *n* = 11 cells, 481 sweeps; difopein *n* = 9 cells, 448 sweeps) revealed that multi-channel openings were more likely in 14-3-3ε overexpressing cells (Fig. 1 J-L). In contrast, we observed reduced channel availability (likelihood of observing at least one event/sweep) in difopein expressing cells (Fig. 1 N). With our results supporting a role for 14-3-3 in promoting surface expression and availability of Ca_V_1.2 channels, we reasoned this could have implications for their cooperative gating. This gating modality occurs when Ca_V_1.2 channels physically interact within clusters, where they influence each other’s activity so that the opening of one channel increases the likelihood of opening of the attached neighboring channels resulting in an amplification of Ca^2+^ influx (37–39). Overexpression of 14-3-3ε significantly increased the slope steepness of Boltzmann function used to fit the G/G_max_ data compared to control while a shallower slope was observed in the difopein group (*SI appendix* Table S1) suggesting 14-3-3ε increased cooperativity of the channels while difopein decreased it. To further quantify cooperativity within our *i*_Ca_ data, we applied a coupled Markov chain model (37, 39–42). This model assigns a cooperativity coefficient (κ) to each sweep with a value of 0 indicating exclusively independently gating channels and a value of 1 indicating exclusively cooperatively gating channels underlying the currents. 14-3-3ε overexpressing cells had a trending increase in κ (Fig. 1 O), while difopein expressing cells displayed a slight decrease in κ (Fig. 1 O) compared to control cells. Taken together these data suggest a possible role for 14-3-3ε in promoting cooperative interactions between Ca_V_1.2 channels.

### 14-3-3 cannot drive surface trafficking of Ca_V_α_1C_ without Ca_V_β

Given the clear Ca_V_1.2 trafficking enhancing effects of 14-3-3ε we wondered if 14-3-3ε overexpression could drive trafficking of the Ca_V_α_1C_ subunit to the plasma membrane without a requirement for co-expression of Ca_V_β auxiliary subunits. Indeed a prior study found 14-3-3τ overexpression sufficient to force trafficking of Ca_V_α_1B_ (pore-forming subunit of Ca_V_2.2) in tsA-201 cells without co-expression of any auxiliary subunits (28); to test if a similar mechanism exists for Ca_V_1.2, we overexpressed 14-3-3ε with Ca_V_α_1C_ and Ca_V_α_2_δ without the Ca_V_β subunit and found that 14-3-3ε overexpression was not sufficient to drive trafficking of Ca_V_1.2 independent of the Ca_V_β subunit (*SI appendix* Fig. S2). Rather, 14-3-3ε appears to play a co-regulatory role in Ca_V_α_1C_ trafficking since its overexpression enhanced channel surface expression only in the presence of Ca_V_β.

### β-AR stimulation enhances 14-3-3 – Ca_V_1.2 colocalization on the t-tubule sarcolemma

Given our finding that endogenous 14-3-3 co-IPs with Ca_V_1.2 in tsA-201 cell lysates, we next investigated the relative localization of the two proteins in isolated ventricular myocytes. Although 14-3-3 is a cytosolic protein, it is known to associate with many membrane proteins and has been shown to accumulate in areas where it has strong regulatory effects such as synapses (43) and intercalated disks (20). We examined 14-3-3 and Ca_V_1.2 distribution and colocalization in freshly isolated adult mouse ventricular myocytes (AMVMs) using Airyscan super-resolution imaging of cells immunolabeled for Ca_V_1.2 and 14-3-3 in control and ISO-stimulated conditions (Fig. 2A). In methanol and Triton X-100 permeabilized cells, much of the cytosolic 14-3-3 is lost but membrane protein-bound 14-3-3 appeared to be retained, and a striking z-line 14-3-3 localization pattern was evident (Fig. 2A). A similar distribution was observed in paraformaldehyde-fixed cells permeabilized with either saponin or Triton X-100 (*SI appendix* Fig. S3). In unstimulated (no ISO) control myocytes, 20.92 ± 1.99 % of the endogenous 14-3-3 was found colocalized with Ca_V_1.2 at this resolution (Fig. 2A-B). Acute ISO (100 nM for 8 mins) stimulation of isolated ventricular myocytes prior to fixation resulted in a significant increase in 14-3-3 – Ca_V_1.2 colocalization to 27.50 ± 2.26 % (Fig. 2A-B). This ISO-stimulated increase in 14-3-3 – Ca_V_1.2 interactions was further confirmed using proximity ligation assays (PLA; Fig. 2C-D). In this technique, ventricular myocytes were immunostained with anti-Ca_V_1.2 and anti-14-3-3 primary antibodies coupled to species-specific DNA primer tagged secondary antibodies. When the two sets of primer-tagged probes (Ca_V_1.2-PLUS and 14-3-3-MINUS) come within 40 nm of one another, they hybridize. Following ligation to form circular DNA, amplification of the DNA and addition of a fluorescent detection reagent allows visualization of fluorescent puncta as a readout at the sites of proximity (44). Accordingly, we observed 23.35 % more puncta on average in ISO-stimulated myocytes compared to unstimulated controls (Fig. 2D). Together with the Airyscan data, these results suggest that 14-3-3 associates with Ca_V_1.2 channels on the t-tubule sarcolemma and that the association is enhanced upon activation of the *β*-AR signaling cascade.

**Figure 2.**
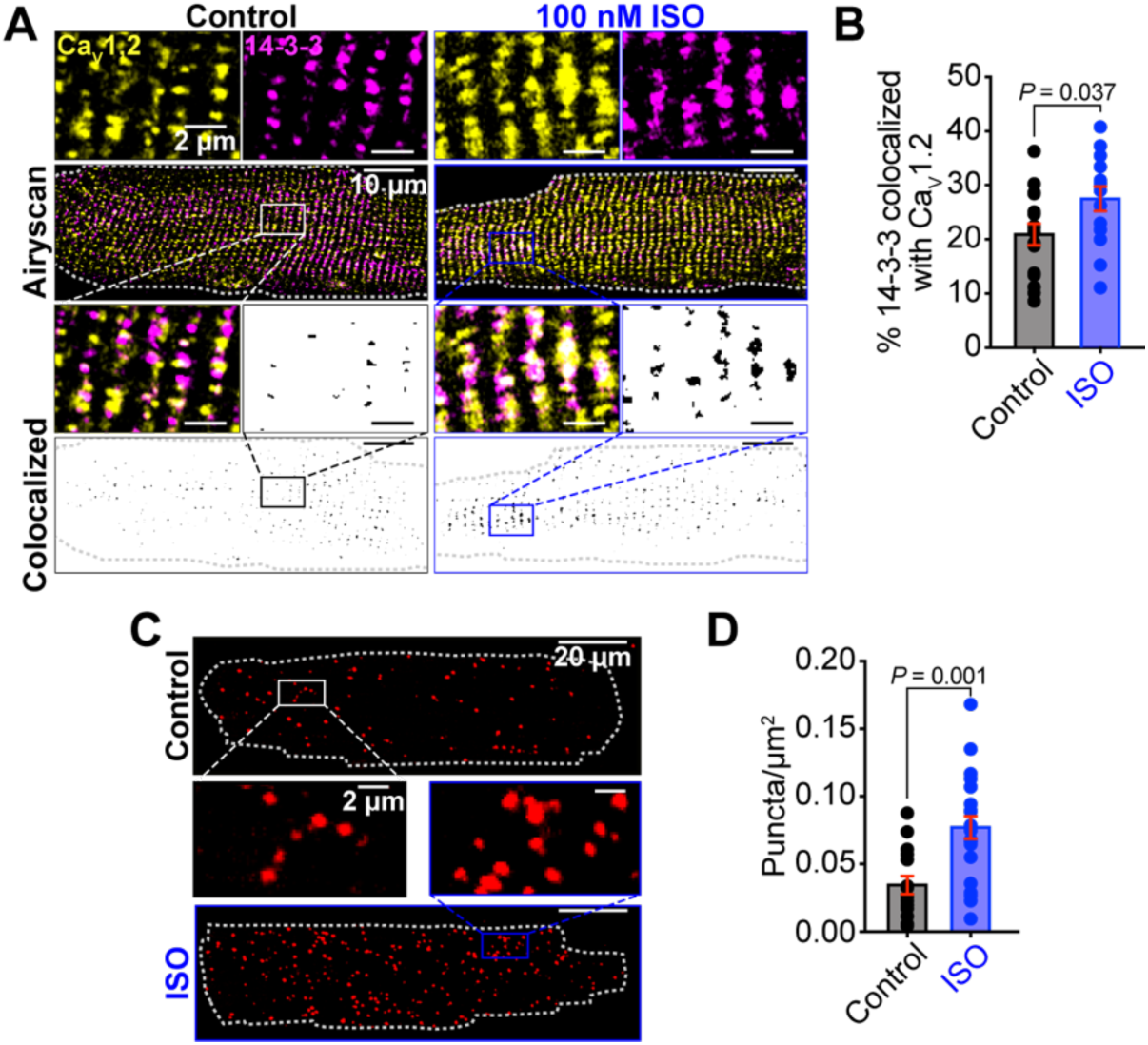
14-3-3 and Ca_V_1.2 colocalize in cardiomyocytes, and isoproterenol stimulates an increase in this colocalization. **A**, Airyscan images showing Ca_V_1.2 and 14-3-3 localization in representative immunostained myocytes without (left) and with (right) ISO. *Bottom*: Binary colocalization maps showing regions of pixel overlap between signals. **B**, histogram summarizing % colocalization between 14-3-3 and Ca_V_1.2 in myocytes in control and ISO-stimulated conditions (control: *N* = 4, *n* = 16; ISO: *N* = 3, *n* = 14). **C**, representative PLA images showing proximity between 14-3-3 and Ca_V_1.2 with and without ISO. **D**, quantification of the proximity sites (puncta) per μm^2^ in myocytes (control: *N* = 3, *n* = 15; ISO: *N* = 3, *n* = 22). Data were analyzed using unpaired Student’s t-tests. Error bars indicate SEM.

### 14-3-3 interactions affect the nanoscale distribution and clustering of Ca_V_1.2 channels

We have previously reported that Ca_V_1.2 recycling from endosomal reservoirs is enhanced by acute *β-*AR stimulation with ISO (45–48). This enhanced recycling leads to mobilization of endosomal Ca_V_1.2 channels to the t-tubule sarcolemma where they form larger clusters. Thus, we hypothesized that the increase in colocalization between 14-3-3 and Ca_V_1.2 with ISO could simply occur because of that increased t-tubular localization and clustering of Ca_V_1.2. We tested this hypothesis by performing single molecule localization microscopy (SMLM; lateral resolution of ∼30 nm) to examine the nanoscale distribution, clustering, and colocalization of 14-3-3 and Ca_V_1.2 in control and ISO-stimulated myocytes. At this enhanced resolution the z-line pattern of 14-3-3 and Ca_V_1.2 localization remained evident and discrete areas of co-localization of the two proteins were observed as in the Airyscan imaging experiments. Consistent with previous findings, Ca_V_1.2 clustering and expression along the t-tubule sarcolemma significantly increased (Fig. 3A-C). Although we expected 14-3-3 cluster size and expression to concurrently increase, neither parameter was significantly increased after ISO (Fig. 3A-C). Rather than co-trafficking with the channel, this suggests that 14-3-3 may be acting as an organizing point or nucleation factor for Ca_V_1.2 super-clustering. In these experiments, we also saw a significant increase in the percent of 14-3-3 colocalized with Ca_V_1.2, but not in the percent of Ca_V_1.2 colocalized with 14-3-3 (Fig. 3D-E). Accordingly, we measured the Ca_V_1.2 cluster size versus their distance to 14-3-3 and found that not only were the clusters in contact with 14-3-3 larger in control conditions, only this population increased after ISO (Fig. 3F-H). Collectively these results suggest that ISO-stimulated recycling of Ca_V_1.2 channels selectively occurs at sites on the t-tubule sarcolemma where 14-3-3 is already present.

**Figure 3.**
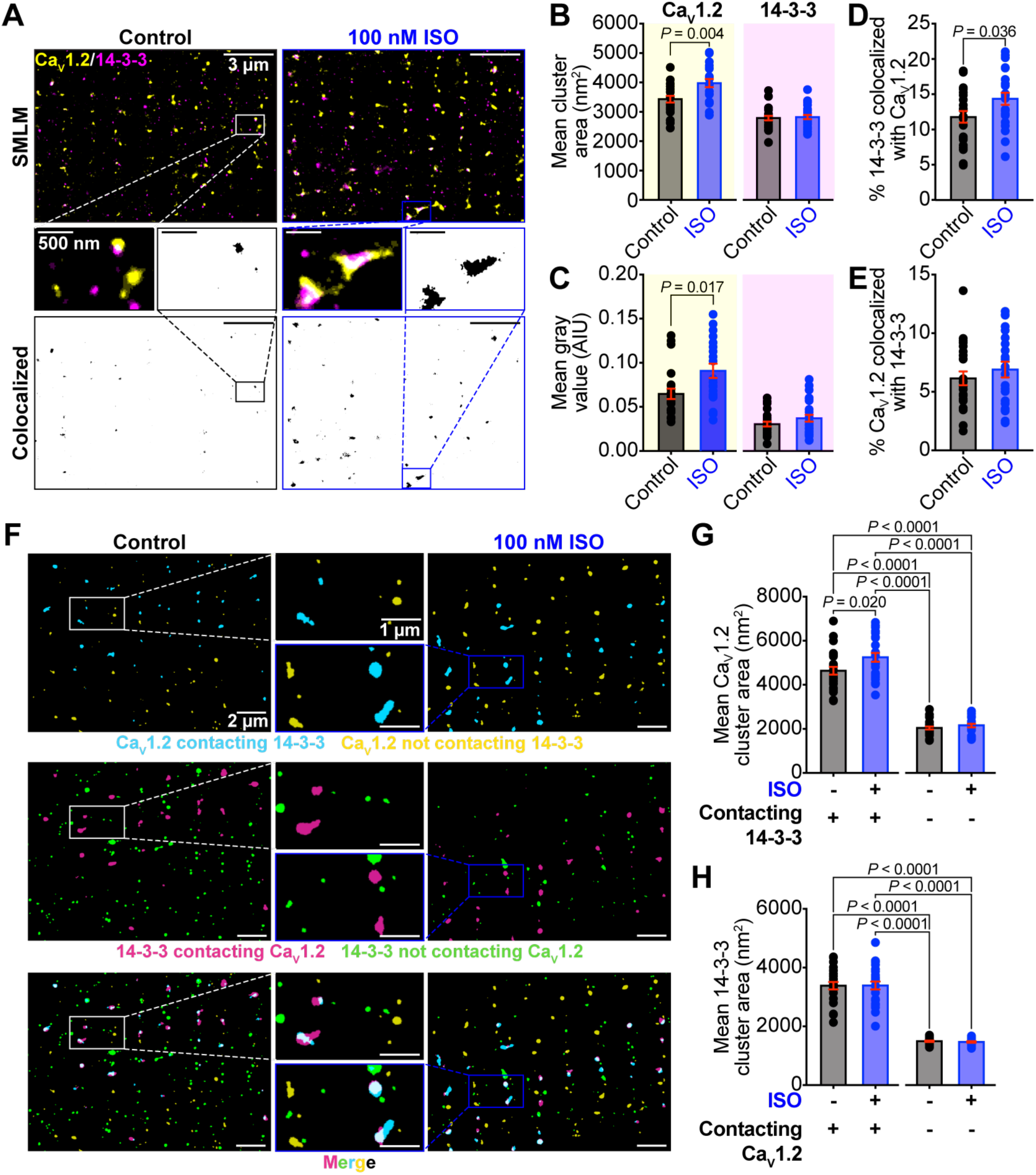
14-3-3 acts as a nucleation factor for *β*-AR stimulated Ca_V_1.2 super-clustering. **A,** SMLM maps of cardiomyocytes immunostained against Ca_V_1.2 and 14-3-3 in myocytes with or without ISO-stimulation. **B-C**, histograms summarizing mean Ca_V_1.2 channel cluster area (control: *N* = 3, *n* = 21; ISO: *N* = 3, *n* = 20), 14-3-3 channel cluster area (control: *N* = 4, *n* = 25; ISO: *N* = 4, *n* = 25), Ca_V_1.2 gray value (control: *N* = 4, *n* = 23; ISO: *N* = 4, *n* = 22), and 14-3-3 gray value (control: *N* = 4, *n* = 25; ISO: *N* = 4, *n* = 25). **D**, histograms summarizing % colocalization between 14-3-3 and Ca_V_1.2, and **E**, % colocalization between Ca_V_1.2 and 14-3-3 in myocytes with and without ISO (control: *N* = 4, *n* = 24; ISO: *N* = 4, *n* = 22). **F,** SMLM maps showing 14-3-3 clusters in contact (magenta) and not in contact (green) with Ca_V_1.2, and Ca_V_1.2 clusters in contact (teal) and not in contact (yellow) with 14-3-3. **G,** histogram summarizing mean Ca_V_1.2 channel cluster area (control: *N* = 4, *n* = 25; ISO: *N* = 4, *n* = 24) and **H,** 14-3-3 channel cluster area (control: *N* = 3, *n* = 24; ISO: *N* = 4, *n* = 24). Data were analyzed using unpaired Student’s t-tests (B, D, and E), and Mann Whitney tests (C). Data in G and H were analyzed using two-way ANOVAs with multiple comparison post-hoc tests. Error bars indicate SEM.

### Phosphorylation-dependent 14-3-3 binding sites on Ca_V_1.2

With accumulating evidence pointing toward phosphorylation augmented 14-3-3 binding to Ca_V_1.2, we next mapped potential binding sites for 14-3-3 on Ca_V_1.2 by overlaying known phosphorylation sites (49) with the 14-3-3 binding motifs using 14-3-3-Pred, a web-based tool that predicts 14-3-3 binding sites in proteins (33) (*SI appendix* Fig. S4A). The rabbit gene was used for numbering for Ca_V_α_1C_ and Ca_V_β_2b_, but equivalent sites also exist for the human and mouse genes. 14-3-3 dimers have two amphipathic binding grooves that can permit binding of pairs of phosphorylated residues located >15 amino acids apart (32). Two optimal consensus binding motifs for 14-3-3 have been identified, mode I R(S/X)X(pS/T)XP, and Mode II RX(F/Y)X(pS/T)XP where X is any amino acid and pS/T denotes phosphorylated serine or threonine (32). An additional third mode allows a lower affinity binding of 14-3-3 to the final few amino acids at a protein C-terminal end accordingly the mode III motif is p(S/T)X_1−2_-COOH (50). However, several non-canonical 14-3-3 binding motifs have been reported that can subtly or even quite wildly deviate from these, including some that do not require phosphorylation (51). There are many phosphorylation sites on both the Ca_V_α_1C_ pore-forming subunit of Ca_V_1.2 and the auxiliary Ca_V_β_2b_ subunit (49), but only five along the C-terminal tail of the Ca_V_α_1C_ subunit, and three in the Ca_V_β_2b_ subunit were identified as being putative 14-3-3 binding sites. Of interest, an additional 14-3-3 binding site was identified at the extreme distal C-terminus of the Ca_V_α_1C_ subunit which allows for the possibility of mode III binding, although this serine is not a known phosphorylation site. In this binding mode only one of the 14-3-3 dimer protein binding pockets is occupied, making the interaction with the client protein unstable. These interactions can be dramatically stabilized by the fungal toxin fusicoccin A (FC-A) which occupies the vacant binding pocket in the 14-3-3 dimer to stabilize the interaction (52). We reasoned that addition of FC-A to our assay would allow us to resolve unstable mode III interactions if they were present between 14-3-3 and Ca_V_1.2. If the putative 14-3-3 biding site on the extreme distal C-terminal is significantly involved in the phosphorylation-dependent binding of 14-3-3 to Ca_V_1.2 then we predicted that stabilization of that interaction with FC-A would result in a more prominent FSK-stimulated interaction in our co-IPs. To test that, we performed co-IPs in the presence of FC-A, however we observed no appreciable difference between FC-A treated and untreated samples, with no additional FSK effects (*SI appendix* Fig. S4B). These data suggest that this Ca_V_α_1C_ distal C-terminal site is not a major binding site for 14-3-3 to Ca_V_1.2 but it remains to be determined which of the other identified sites are essential for these interactions.

### 14-3-3 and Ca_V_1.2 interactions with Rad

Considering the known interactions between 14-3-3 and the small RGK protein Rad (30, 31), and between Rad and the Ca_V_β subunit of Ca_V_1.2 (8–12), we investigated the dependence of 14-3-3 and Ca_V_1.2 interactions on the presence of Rad. If both sets of proteins interact with each other, the interaction we saw could be due to formation of a larger complex and not direct binding. Therefore, we probed for GFP-tagged Rad in tsA-201 cell co-IPs with and without Rad transfected. Rad contains two paired phosphorylation sites required for 14-3-3 binding, suggesting both sides of the dimer bind Rad simultaneously (31)( *SI appendix* Fig. S4A). Both of these sites are known PKA-phosphorylation sites (8), and indeed we saw an increase in the amount of Rad in the 14-3-3 IP with FSK treatment in cells co-transfected with Rad, confirming that our FSK treatment was successful (*SI appendix* Fig. S4C). Interestingly, in our hands the GFP-tagged Rad did not pull down with Ca_V_1.2 in any condition. Given that we did not see Rad and Ca_V_1.2 binding at all, and that the degree of association between Ca_V_1.2 and 14-3-3 was similar in cells with or without Rad co-transfection (data not shown) we can be confident that the Ca_V_1.2 ***–*** 14-3-3 interactions that we observed were not dependent on Rad and instead imply direct binding between 14-3-3 and Ca_V_1.2.

### Modeling of Ca_V_1.2 interaction with 14-3-3 using AlphaFold 3

Seeking insight into possible interaction sites between Ca_V_1.2 and 14-3-3ε, we utilized AlphaFold 3 (AF3) (53) to model the interaction between the α_1C_ and β_2_ subunits of Caᵥ1.2 and a dimer of 14-3-3ε, including phosphorylation in the C-terminal region of the α subunit as described in Methods. AF3 is a diffusion-based deep neural network capable of predicting the biological structure of protein complexes with nucleic acids, small molecules, ions, and post-translational modifications, including protein phosphorylation. The AF3-predicted model of rabbit and human Caᵥ1.2 closely resembles cryo-electron microscopy (cryoEM) structure of human Caᵥ1.2 (PDB: 8we6) (54), with a root mean square deviation (RMSD) between Cα atoms of ∼2.8 Å and ∼3.2 Å, respectively, over the experimentally resolved regions of the human Caᵥ1.2 structure (*SI appendix* Fig. S5A). Similarly, the AF3 model of a human 14-3-3ε dimer closely agreed with the published x-ray structure (PDB: 7c8e) (55), with RMSD ∼0.9 Å (*SI appendix* Fig. S5B). AF3 per-residue predicted local distance difference test (pLDDT) confidence scores in the rabbit Caᵥ1.2 model of the C-terminal region varied from very low (21) to low (50–70) to good (72) levels (*SI appendix* Fig. S6), corresponding to the comparatively less ordered nature of the C-terminal region which has so far not been fully resolved. The model failed to predict any association between Caᵥ1.2 α_1C_ and 14-3-3ε in the absence of phosphorylation, whereas the addition of phosphate groups at S1700 and S1928 allowed interactions of Caᵥα_1C_ with two 14-3-3ε molecules, in the conformation expected of a dimer (*SI appendix* Fig. S5C-D). Notably, pLDDT confidence in the C-terminal regions containing phosphorylation sites was very low (41–46) at pS1700 and pS1928 in the AF3 model of rabbit Caᵥ1.2, and low (∼52) at pS1718 and pS1981 in the AF3 model of human Caᵥ1.2 (*SI appendix* Fig. S6). Despite these limitations this AF3-generated model predicts a phosphorylation-dependent association between a dimer of 14-3-3ε and the C-terminal tail of Ca_V_1.2 α_1C_ via pS1928 and pS1700, two residues known to be phosphorylated during *β*-AR stimulation *in vivo*.

### Acute 14-3-3 inhibition impairs ISO-stimulated Ca_V_1.2 channel recycling

We hypothesized that a phosphorylation dependent association between 14-3-3 and Ca_V_1.2 in cardiomyocytes that impacts ISO-stimulated channel super-clustering would have an impact on the functional responsivity and redistribution of Ca_V_1.2 upon *β*-AR activation. To begin addressing that we first tested the role of endogenous 14-3-3 in basal Ca_V_1.2 channel clustering and ISO-stimulated superclustering, using ventricular myocytes acutely treated with the cell-permeable 14-3-3 inhibitor BV02 (50 µM, for 8 mins, at 37 °C). Cells were subsequently immunostained and imaged using SMLM. Results revealed that acute BV02 treatment did not significantly affect basal cluster size, localization, or expression of Ca_V_1.2 compared to PBS-treated controls however it significantly attenuated the ISO-stimulated superclustering response (Fig. 4A-C). These data support a model where 14-3-3 plays a critical role in ISO-stimulated Ca_V_1.2 endosomal recycling and/or insertion of channels into the membrane but does not overtly affect the basal sarcolemmal Ca_V_1.2 population at least on this acute timescale.

**Figure 4.**
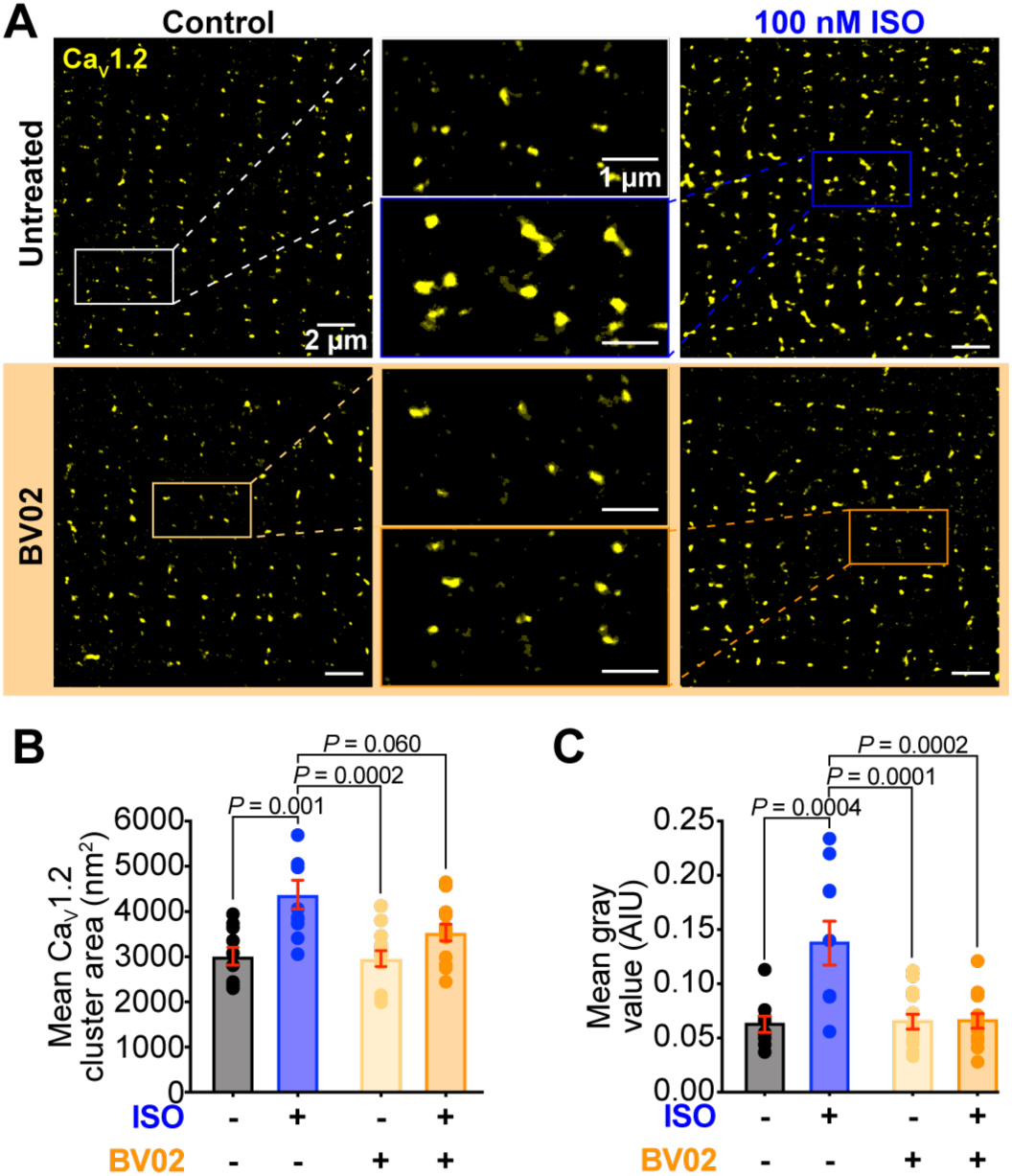
Acute 14-3-3 inhibition abrogates *β*-AR stimulated Ca_V_1.2 super-clustering in isolated cardiomyocytes. **A,** Representative SMLM localization maps of cardiomyocytes immunostained against Ca_V_1.2 in untreated, 100 nM ISO-treated, 50 μM cell permeable 14-3-3 inhibitor BV02-treated, and combined ISO and BV02-treated conditions. **B,** histograms summarizing mean Ca_V_1.2 channel cluster area (*N* = 3 for all groups, control: *n* = 10; ISO: *n* = 10; BV02: *n* = 14; ISO-BV02: *n* = 13), and **C,** Ca_V_1.2 gray value (*N* = 3 for all groups, control: *n* = 9; ISO: *n* = 10; BV02: *n* = 14; ISO-BV02: *n* = 13). Data were analyzed using two-way ANOVAs with multiple comparison post-hoc tests. Error bars indicate SEM.

### 14-3-3 overexpression or prolonged inhibition alters Ca_V_1.2 responsivity to β-adrenergic stimulation in cardiomyocytes

We next examined the effects of more prolonged alterations in 14-3-3 in isolated cardiomyocytes. Cells were cultured for up to 48 hrs to allow adenoviral transduction with either red fluorescent protein (RFP; control), 14-3-3ε-mRuby, or difopein-YFP. Subsequent SMLM experiments were performed on unstimulated and ISO-stimulated cohorts of cells immunolabeled against Ca_V_1.2 and 14-3-3. SMLM performed on cultured cells transduced with RFP confirmed that a robust and significant ISO-stimulated Ca_V_1.2 superclustering response persists in cultured myocytes despite an overall reduction in Ca_V_1.2 channel cluster size and expression compared to uncultured cells (compare Fig. 5A, D and E, to Fig. 3A-C). Whole cell patch clamp recordings of the RFP transduced control cells further confirmed that *β*-adrenergic regulation of *I*_Ca_ was present in these cells with ISO-eliciting a 1.73 ± 0.21-fold increase in peak current density (Fig. 5F-H), and a significant leftward-shift and steeper slope in the voltage dependence of conductance (Fig. 5I, *SI appendix* Table S2) as expected. Overexpression of 14-3-3ε produced a trending increase in basal Ca_V_1.2 channel cluster size and expression that failed to reach significance in a two-way ANOVA compared to RFP-transduced controls (Fig. 5B, D, and E) and produced a similar trend toward an increased basal current density (Fig. 5F and G) while the ISO-stimulated superclustering, and current regulation effects remained intact (Fig. 5B, and D-I). Difopein-mediated 14-3-3 inhibition also had no significant impact on basal Ca_V_1.2 channel clustering or expression but in agreement with the acute BV02-mediated inhibition results, difopein-transduced cells failed to display an ISO-stimulated superclustering response (Fig. 5C-E). Furthermore, while basal Ca_V_1.2 current density was similar to RFP-transduced controls, difopein-transduced cells displayed a significantly reduced responsivity to ISO displaying only a 1.13 ± 0.10-fold increase in peak current density (Fig. 5 F-H). Interestingly, difopein-transduced myocytes continued to display an ISO-stimulated leftward-shift with a similar V_1/2_ despite the loss of peak current enhancement and channel superclustering (*SI appendix* Table S2). These data reveal that 14-3-3 plays a critical role in the ISO-stimulated recycling and superclustering of Ca_V_1.2 channels in cardiomyocytes and further suggest that the enhanced *I*_Ca_ density associated with *β*-adrenergic regulation of Ca_V_1.2 is blunted when 14-3-3 is inhibited.

**Figure 5.**
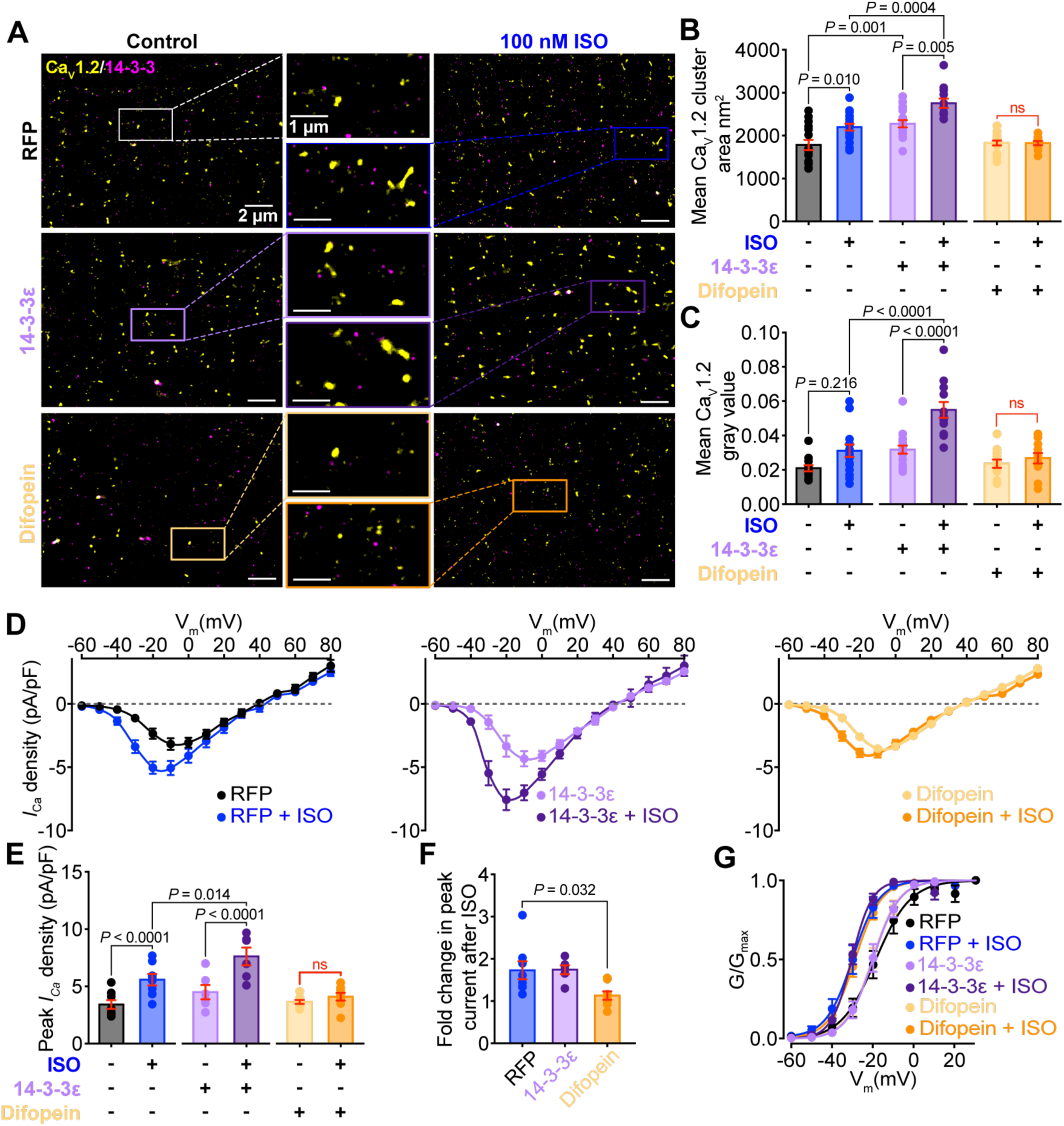
14-3-3 overexpression enhances basal Ca_V_1.2 density, cluster size, and *I_Ca_*, with a preserved *β*-AR response, while 14-3-3 inhibition prevents the *β*-AR stimulated superclustering and increased *I_Ca_* response. **A**, Representative SMLM maps showing Ca_V_1.2 and 14-3-3 localization with and without ISO-treatment in immunostained cardiomyocytes cultured for 48 hrs with Ad-RFP (*control*), Ad-14-3-3ε-mRuby, or Ad-difopein-EYFP. **B-C,** histograms summarizing mean Ca_V_1.2 channel cluster area (*N* = 4 for all groups, RFP control: *n* = 15, ISO: *n* = 18; 14-3-3ε control: *n* = 17, ISO: *n* = 12; difopein control: *n* = 16, ISO: *n* = 12), and Ca_V_1.2 gray area (*N* = 4 for all groups, RFP control: *n* = 12, ISO: *n* = 17; 14-3-3ε control: *n* = 17, ISO: *n* = 12; difopein control: *n* = 16, ISO: *n* = 11). **D**, I-V plots and **E,** histogram summarizing peak *I_Ca_* density (control RFP: *N* = 6, *n* = 8; ISO RFP: *N* = 6, *n* = 8; Control 14-3-3ε: *N* = 5, *n* = 6; ISO 14-3-3ε: *N* = 5, *n* = 6; control difopein: *N* = 5, *n* = 8; ISO difopein: *N* = 5, *n* = 8). **F**, fold change in peak *I_Ca_* density following ISO treatment of the cells in D. **G**, voltage-dependence of G/G_max_ fit with Boltzmann functions for cells shown in D. Data were analyzed using two-way ANOVAs (B, C, and E), or one-way ANOVAs (F) with multiple comparison post-hoc tests. Error bars indicate SEM.

## Discussion

We report five lines of evidence supporting a novel regulatory mechanism for Ca_V_1.2 via interactions with 14-3-3ε: 1) co-IPs showed Ca_V_1.2 forms complexes with 14-3-3 in tsA-201 cells; 2) in adult mouse cardiomyocytes Airyscan imaging and PLA revealed 14-3-3 is localized alongside Ca_V_1.2 in the t-tubule sarcolemma and the association between them is increased following *β*-adrenergic stimulation; 3) super-resolution imaging revealed 14-3-3 interactions affect the nanoscale distribution and clustering of Ca_V_1.2 during *β*-adrenergic signaling; 4) examination of the basal Ca_V_1.2 surface expression, whole cell calcium current, single calcium channel currents, and nanoscale distribution of Ca_V_1.2 with overexpressed or inhibited 14-3-3ε revealed 14-3-3 facilitates trafficking and stimulated recycling of Ca_V_1.2; and 5) patch clamp and super-resolution imaging revealed prolonged inhibition of 14-3-3 or overexpression of 14-3-3ε alters Ca_V_1.2 plasma membrane localization, basal function, and *β*-adrenergic responsivity.

14-3-3 is known to interact with many ion channels and has recently been appreciated as an important regulator of many excitation-contraction (EC) coupling proteins. In the heart, 14-3-3 regulates Na_V_1.5 (20, 21), K^+^ channels such as HERG (18), TASK-1 (19) and K_ATP_ (15), and Ca^2+^ handling proteins such as SERCA through phospholamban (23), PMCA (24, 25) and NCX (26). In the brain and in heterologous expression systems, 14-3-3 also regulates voltage-gated N-type Ca^2+^ channels (Ca_V_2.2), a structurally similar calcium channel involved in neurotransmitter release (27, 28). Despite these discoveries, there has been no direct investigation of Ca_V_1.2 regulation by 14-3-3. A study pursuing novel approaches to inhibit calcium channels found that artificially membrane-bound 14-3-3ε was able to inhibit Ca_V_1.2, presumably by binding to the channel and pulling a cytosolic portion of the channel into an unfavorable position (29). Although this study showed an interaction between the modified 14-3-3ε and Ca_V_1.2, the endogenous effects of 14-3-3 on Ca_V_1.2 were not examined. Our study provides the first evidence for complex formation between 14-3-3 and Ca_V_1.2, both in heterologous systems and cardiac myocytes.

Our results support a role for 14-3-3 in regulation of Ca_V_1.2 channel expression at the plasma membrane. In tsA-201 cells, 14-3-3ε overexpression promoted enhanced expression of Ca_V_1.2 at the plasma membrane while competitive inhibition of 14-3-3 had the opposite effect, resulting in a near total inability of the channels to get to the membrane. This dependence of Ca_V_1.2 trafficking on 14-3-3 in tsA-201 cells is supported by both imaging and electrophysiology experiments in which *I*_Ca_ and single channel activity were extremely low in cells transfected with difopein compared to controls transfected with or without inactive difopein. In that series of experiments, it was difficult to determine whether a patched cell displayed no current because some element of the channel complex had not transfected adequately or because of an effect of difopein thus a criterion for inclusion in the analysis was that the cell must have some measurable Ca^2+^ current. This level of caution likely means we are underestimating the effect of 14-3-3 inhibition. Our findings are consistent with the report that 14-3-3τ promotes plasma membrane localization of voltage-gated N-type Ca_V_2.2 Ca^2+^ channels in tsA-201 cells (28). Although tsA-201 cells endogenously express all transcripts of 14-3-3 isoforms (56), our results indicate that endogenous or even overexpressed 14-3-3 is not sufficient to drive membrane expression of Ca_V_1.2 channels in the absence of auxiliary subunits, unlike for Ca_V_2.2. Rather it seems that 14-3-3 acts in concert with the auxiliary subunits to promote membrane expression of Ca_V_1.2 α_1C_. It is generally accepted that Ca_V_β subunits are required for trafficking of α_1C_ however a recent study reported that cardiac α_1C_ can traffic to the sarcolemma of cardiomyocytes even when they have a mutation that abrogates their binding to Ca_V_β subunits (57). While interpretation of those results is complicated by the fact that the mutant channel trafficking was studied in cells that also expressed WT channels that could bind endogenously expressed Ca_V_β subunits, it remains an untested possibility that endogenous 14-3-3 could facilitate forward trafficking of those mutant channels in cardiomyocytes.

One of the original hypotheses as to how Ca_V_β could facilitate Ca_V_1.2, trafficking was that its binding to the AID region of the I-II loop of the channel masked an unidentified ER-retention motif (58). That motif was never identified and the hypothesis was largely rejected (59, 60) with subsequent work revealing that the proximal C-terminus of Ca_V_1.2, far from the AID, is critical for membrane targeting of α_1C_ (61). CD4-fusion constructs of various segments of the Ca_V_1.2 α_1C_ revealed two putative ER-retention motifs in the proximal C-terminal of Ca_V_1.2, and one in a similar region of Ca_V_2.2 (60). A similar approach using Myc-CD8α fused to segments of the C-terminal of rat Ca_V_2.2 confirmed that amino acids 1706-1940 on the proximal C-terminal contains an ER-retention signal that can be masked by 14-3-3 binding to allow surface expression of the channel (28). A second phospho-regulated 14-3-3 binding site on the distal Ca_V_2.2 C-terminal (S2126) was found to affect the inactivation properties of the channel, although they did not observe changes to Ca_V_1.2 inactivation in the same conditions (27, 28). In line with these results, we did not observe any effects of 14-3-3 on Ca_V_1.2 inactivation properties, however mapping of the possible 14-3-3 binding sites predicted five phosphorylation-dependent 14-3-3 binding sites on the C-terminal tail of Ca_V_1.2, in addition to the C-terminal site. Of these five, two are in similar positions to the sites found on Ca_V_2.2, one located proximally (S1700), and another located distally (S1928). Interestingly, mutant animals with phosphorylation-preventing alanine substitutions of S1700 and T1704 on the C-tail of Ca_V_1.2 display significantly suppressed sarcolemmal expression of the channel and a correspondingly reduced basal *I*_Ca_ but intact responsivity to *β*-adrenergic receptor stimulation (62). A similar mutant mouse generated independently displayed a similarly reduced basal *I*_Ca_ but was said to have similar expression levels of the channel, however a definitive biotinylation-assay was not performed to quantify the membrane population (63). It is possible that 14-3-3 binding to phosphorylated S1700 or perhaps T1704 masks an ER-retention signal located in that proximal segment of the C-terminal tail, facilitating surface expression of the channels. In contrast, phosphorylation of the S1928 site has been associated with PKA-dependent acute increases in Ca_V_1.2 activity in vascular smooth muscle (64) and neurons (65). Although this site is a PKA target in cardiac Ca_V_1.2 as well, there has been no such definitive regulatory role for S1928 phosphorylation in the heart (66). Binding of 14-3-3 to both sites simultaneously may allow phosphorylation-dependent trafficking of Ca_V_1.2 based on the channel’s proximity to *β*-adrenergic signaling complexes. AF3 modeling confirmed a potential structure of Ca_V_1.2 with 14-3-3 bound concurrently at S1700 and S1928.

Our findings imply that the surface expression of Ca_V_1.2 is promoted by their phosphorylation-dependent association with 14-3-3. Phosphorylation-dependent anterograde trafficking of ion channels has been proposed for several K^+^ channels (15, 19, 67), Na^+^ channels (68), and Ca_V_1.2 Ca^2+^ channels (62, 69). In the case of TASK potassium channels the underlying mechanism has been well-studied and involves phosphorylation and 14-3-3-binding acting as a “release” switch that prevents an otherwise constitutive binding of the channels to coat protein (COP) complex I (COPI) (19, 70). COPI mediates retrograde transport of proteins from endosomes back to the Golgi and from the Golgi back to the ER but phosphorylation of TASK-3 and binding of 14-3-3 has been proposed to release the channels from COPI to allow their anterograde transport and membrane expression. Cardiac ATP-sensitive potassium channels have been found to display a similar increase in sarcolemmal expression in response to their phosphorylation downstream of *β*-AR stimulation (15). Furthermore, previous work from our group revealed that endosomal reservoirs of Ca_V_1.2 channels are recycled back to the membrane in a phosphorylation-dependent manner upon *β*-adrenergic receptor stimulation (45–48). These prior results from our group and others, paired with the novel findings in the present study, and our AF3-predicted model of association between Ca_V_1.2 and 14-3-3ε invite the speculative conclusion that a similar mechanism could be in place for these voltage-gated Ca^2+^ channels.

In cardiomyocytes inhibition of 14-3-3 with either difopein or BV02 did not overtly alter peak *I*_Ca_ or the basal expression of the channels in-so-far as we can tell from imaging experiments. Since both difopein and BV02 are competitive inhibitors of 14-3-3 it may be that we are not achieving high enough concentrations of these agents to have the desired dominant negative effect. It may also be a case of bad timing in that channels may require 14-3-3 to bind to them during their transit through the biosynthetic delivery pathway, thus the effects may be easier to observe on the blank slate of a tsA-201 cell where we can manipulate 14-3-3 levels/binding while the channels are all actively being translated. Thus, effects of 14-3-3 on basal expression may be more difficult to ascertain in ventricular myocytes. However, it was clear that 14-3-3 plays a role in modulating the response to *β*-AR stimulation with 14-3-3 inhibition abrogating the ISO-induced super-clustering of Ca_V_1.2 channels and significantly blunting the *I*_Ca_ augmentation. Interestingly, although the increase in *I_Ca_* was abolished with 14-3-3 inhibition, we still observed a significant left-shift in voltage dependence of activation. This suggests that although both effects are PKA dependent, they are occurring through separate mechanisms, only one of which is regulated by 14-3-3.

The effect of 14-3-3 on ISO-stimulated Ca_V_1.2 channel super-clustering also has implications for cooperative gating of the channels. This gating modality has now been proposed to occur in multiple classes of ion channels where physical interactions between adjacent channels allow them to allosterically communicate such that the opening of one channel in a cluster can increase the probability of gating of the other adherent channels (recently reviewed (38)). We have previously reported that Ca_V_1.2 channels cooperatively gate to enhance calcium influx (37, 39) and this is promoted during *β*-AR activation by the formation of large superclusters of Ca_V_1.2 (45, 46). This process is calmodulin-dependent but based on studies showing 14-3-3 can link Na_V_1.5 channels together, we speculated that 14-3-3 might play an additional role in Ca_V_1.2 cooperative gating, perhaps by linking groups of channels together to form larger oligomers. This function of 14-3-3 in linking proteins into dimers or larger oligomers has been previously observed in both plant and mammalian cells (21, 71). While we did not directly test oligomerization in the present study, we did observe that multi-Ca_V_1.2 channel openings were more likely in 14-3-3ε overexpressing tsA-201 cells, with quantitative measures of cooperativity indicating a positive effect of 14-3-3 and a negative effect of difopein. These suggestive findings will require further study and at present our data do not definitively link 14-3-3 to Ca_V_1.2 oligomerization beyond the observation that it regulates cluster size and channel expression on the plasma membrane.

Based on our data, we propose a novel model in which 14-3-3ε is recognized as a new modulatory protein of Ca_V_1.2 channels, interacting with it in a phosphorylation-dependent manner to regulate the channel’s trafficking, clustering, and responsivity to *β-*AR signaling. Furthermore, we propose that 14-3-3 acts as a nucleation factor for the insertion of Ca_V_1.2 into the sarcolemma during *β*-AR stimulation (Fig. 6). This study adds to the growing list of EC-coupling proteins regulated by 14-3-3 and brings us one step closer to a complete picture of the regulation of Ca_V_1.2.

**Figure 6.**
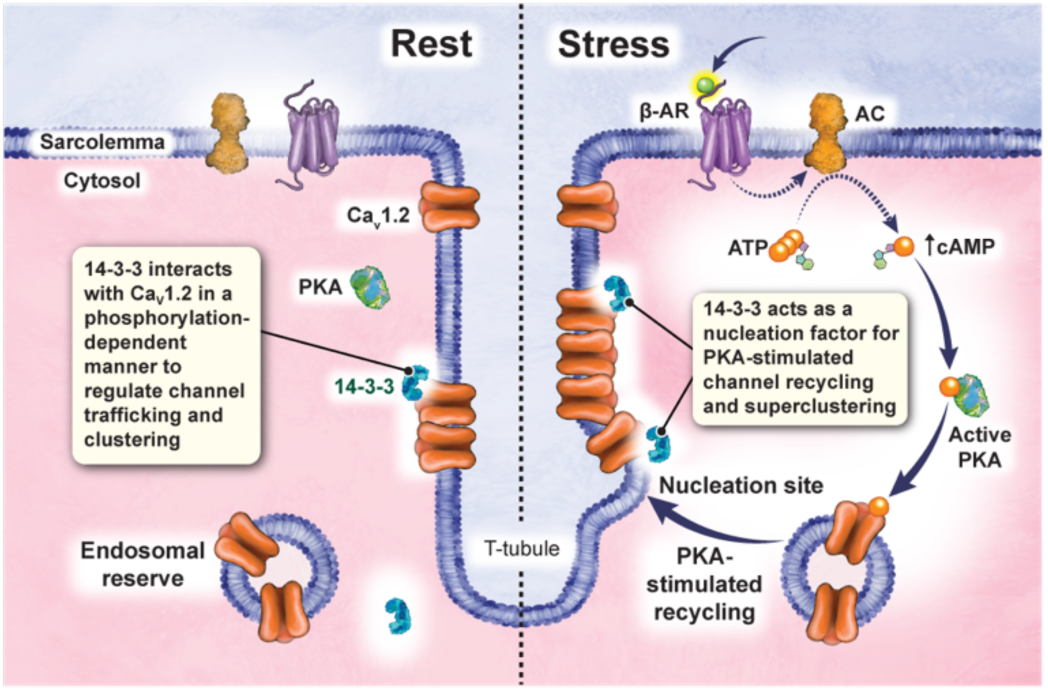
A working model for 14-3-3 regulation of Ca_V_1.2. 14-3-3 binds to Ca_V_1.2 and is localized along the z-lines in cardiomyocytes alongside Ca_V_1.2 channels at rest. This localization can be enhanced during *β*-AR stimulation. During activation of this pathway, 14-3-3 acts as a nucleation factor for the insertion of endosomal Ca_V_1.2 into large super-clusters to dynamically tune Ca_V_1.2 activity and abundance in response to hemodynamic demand.

## Materials and methods

All animal handling and procedures adhered to the National Institutes of Health Guide for the Care and Use of Laboratory Animals (UC Davis) and were approved by the local Institutional Animal Care and Use Committee. tsA-201 cells were transiently transfected with combinations of Ca_V_1.2 α_1c_, Ca_V_β_2b_, Ca_V_α_2_δ, 14-3-3 isoforms, difopein, or inactivated difopein and used for microscopy and patch clamp electrophysiology experiments. Ventricular myocytes were isolated from 3–6-month-old C57BL/6J mouse hearts via retrograde Langendorff perfusion as previously described (45, 46) and used in microscopy and patch clamp electrophysiology experiments. *N* represents the number of animals and *n* represents the number of cells. Data are reported as mean ± SEM. Statistics were performed using Prism (GraphPad Software Inc.), and all data sets were tested for normality. Unpaired Student’s t-tests or Mann Whitney tests were used to compare data sets with two groups, one-way ANOVAs or Krushkal-Wallis (non-parametric) tests with post-hoc testing were used to compare data sets with more than two groups, or two-way ANOVAs when there were two independent variables. *P* < 0.05 was considered statistically significant. Detailed methods can be found in the SI Appendix.

## Data Availability

All data are included in the manuscript and/or SI Appendix.

## Acknowledgments

Illustration in Fig. S4 was generated using Biorender.com. We are grateful to Mr. Joshua Tulman who produced the Graphical Abstract artwork and to Dr. Jody Martin of the UC Davis CVRI who generated the adenoviruses used in this study. This work was supported by NIH grants R01HL159304 and R01AG063796 to R.E.D. and by RF1NS131379 and R35GM149211 to E.J.D.; by T32 GM099608 and F31HL165815 to H.C.S.

## Notes

### Competing Interest Statement

The authors have declared no competing interest.

