## Supplementary Methods and Figures for "14-3-3 promotes sarcolemmal expression of cardiac Ca_V_1.2 and nucleates isoproterenol-triggered channel super-clustering"

**SI Appendix**

**Detailed Materials and Methods**

**Ethical Approval**

University of California Davis Institutional Animal Care and Use Committee (IACUC) approved all procedures involving mice and were in accordance with the Guide for the Care and Use of Laboratory Animals (1). Mice were kept in a vivarium with a standard light-dark cycle and received standard chow (Laboratory Rodent Diet 5001, LabDiet, St Louis, MO, USA) and water *ad libitum*.

**Cardiomyocyte isolation**

12–24-week-old C57BL/6J mice (The Jackson Laboratory, Sacramento, CA, USA) were euthanized using intraperitoneal injection of pentobarbital solution (> 100 mg/kg; B euthanasia-D Special; Merck Animal Health, Madison, NJ, USA). Hearts were then removed with sharp dissection and washed in ice cold digestion buffer (130 mM NaCl [Thermo Fisher Scientific, Waltham, MA, USA], 5 mM KCl [Thermo Fisher Scientific], 3 mM Na-pyruvate [Sigma-Aldrich, St. Louis, MO, USA], 25 mM HEPES [Thermo Fisher Scientific], 0.5 mM $MgCl_2$ [Thermo Fisher Scientific], 0.33 mM $NaH_2PO_4$ [Thermo Fisher Scientific], and 22 mM dextrose [Thermo Fisher Scientific]) supplemented with 150 µM EGTA (Sigma-Aldrich). Ventricular myocytes were then isolated using the Langendorff technique as previously described (2, 3). Briefly, the aorta was cannulated, and the heart was perfused with warm (37°C) digestion buffer containing 150 µM EGTA until the heart was free of blood. Then, the solution was changed to warm digestion buffer with 50 µM $CaCl_2$ (Thermo Fisher Scientific), 0.04 mg/ml protease (Sigma-Aldrich), and 1.4 mg/ml type 2 collagenase (Worthington Biochemical, Lakewood, NJ, USA). When digestion was finished (as assessed by texture and appearance of heart), the atria, aorta, and pulmonary vessels were removed and the ventricular myocytes were separated by pipetting in warm digestion buffer supplemented with 1.2 mg/mL collagenase, 0.04 mg/mL protease, 100 µM $CaCl_2$ and 10 mg/ml BSA (Sigma-Aldrich). After 2 min centrifugation at 300 rpm, supernatant was discarded, and cells were resuspended in room temperature wash buffer made by supplementing digestion buffer with 250 µM $CaCl_2$ and 10 mg/ml BSA. The cells were centrifuged again at 300 rpm, supernatant was discarded, and cells were resuspended in the appropriate solution.

**tsA cell culture and transfection**

tsA-201 cells (Sigma-Aldrich) were maintained in Dulbecco's Modified Eagle's medium (DMEM; Gibco) supplemented with 10% FBS and 1% penicillin/streptomycin in a cell culture incubator

(37°C, 95:5% $O_2$:$CO_2$). Cells were passaged twice a week and used between passages 14-30. Cells were transfected at 70-85% confluency with JetPEI DNA Transfection reagent (Polyplus Transfection, New York, NY, USA) and plated onto glass coverslips at low density after ~30 hrs of transfection, to be used for experiments the following morning ~16 hrs later (total 48 hr transfection). For imaging experiments, coverslips were treated with poly-$_L$-lysine (0.01%) for 20 min then rinsed with DPBS before plating cells.

**Plasmids**

In experiments performed in tsA-201 cells, cells were transfected with 800 ng of the rabbit cardiac isoform of $Ca_V1.2$ $\alpha_{1c}$ (GenBank accession number: NP_001129994.1) and rat $Ca_V\beta_{2b}$ (M88751), and 400 ng rat $Ca_V\alpha_2\delta$ (AF286488). The $Ca_V1.2$ $\alpha_{1c}$ construct was tagged at its C-terminus with a monomeric GFP(A206K) or a tagRFP and the $Ca_V\beta_{2b}$ was tagged with cerulean (a gift from Henry Colecraft). In experiments involving 14-3-3 overexpression, 150 ng of the following 14-3-3 plasmids were used: pcDNA3-HA-14-3-3β (Addgene: 13270), pcDNA3-HA-14-3-3ε (Addgene: 13273), pcDNA3-HA-14-3-3σ (Addgene: 11946), pcDNA3-HA-14-3-3γ (Addgene: 13274), HA-14-3-3η (Addgene: 116887), and HA-14-3-3ζ (Addgene: 116888). For experiments involving difopein inhibition of 14-3-3, 800 ng of the active EYFP-difopein plasmid (pSCM138), or 800 ng of the inactivated EYFP-tagged plasmid (pSCM174) were used (gifts from Dr. Haian Fu; Emory University, Atlanta, GA, USA).

**Co-immunoprecipitation and western blots**

Transfected tsA-201 cells were treated with fresh media or 10 μM forskolin (FSK; Sigma-Aldrich) for 8 min, then rinsed twice with cold PBS with 1 mM $CaCl_2$ and 0.5 mM $MgCl_2$ (PBS-CM), followed by cold $Mg^{2+}$-$Ca^{2+}$-free PBS to lift cells. Cells were centrifuged at 1.5k x g for 10 min at 4°C, resuspended in cold PBS-CM, and centrifuged at 21k x g in a microfuge for 7 min at 4°C. The supernatant was removed, and cells were frozen at -70°C until lysis.

Frozen cell pellets were resuspended in lysis buffer (50 mM Tris-HCL [Thermo Fisher Scientific] pH 7.4, 150 mM NaCl, 5 mM EGTA pH 7.4, 10 mM EDTA [Sigma-Aldrich], and 1% Triton X-100 [Sigma-Aldrich]) with protease inhibitor cocktail (Roche, Basel, Switzerland) and phosphatase inhibitors (1 mM p-Nitrophenyl phosphate [Sigma-Aldrich], 1 mM sodium pyrophosphate [Sigma-Aldrich], 25 mM NaF [Sigma-Aldrich], and 1 μM microcystin LR [Sigma-Aldrich]) and incubated on ice for 10 min. Lysate samples were then centrifuged at 147k x g at 4°C for 1 hr in an ultracentrifuge and the supernatant was kept on ice. Protein A- (Repligen, Boston, MA, USA) or G- (GE Healthcare, Chicago, IL, USA) Sepharose beads were washed in lysis buffer.

Protein concentration was determined by BCA assay (Prometheus Biosciences, Rahway, NJ, USA). Preclear one was made by mixing 10 μL of beads with 10 μg nonspecific (NS) IgG of matching species to the IP antibody (Jackson Laboratory), and lysate. This was incubated on ice for 30 min, then centrifuged at 21k x g in a microfuge at 4°C. The supernatant was transferred to a second tube with the same volume of beads, incubated and centrifuged again for preclear two. This preclear was washed and run on the gel in parallel with the IP lanes as a nonspecific IgG control. The cleared lysate was then transferred to a tube with the same volume of Protein A- (or G-) Sepharose beads and 5 to 10 μg (depending on antibody) primary antibody and left to mix overnight at 4°C. Immunocomplexes bound to the Sepharose beads were harvested by centrifugation at full speed in a microfuge for 5 min at 4°C, then washed in lysis buffer with phosphatase and protease inhibitors, lysis buffer with 500 mM NaCl, and twice in 50 mM Tris-HCl pH 7.4. Co-IP and IgG samples were eluted and denatured in 1.5X reducing Laemmli sample buffer (Invitrogen, Carlsbad, CA, USA) and heated at 70°C for 20 min. Lysate samples were prepared in 1X final concentration reducing Laemmli sample buffer and heated as above.

All lysate samples were loaded on 4-12% Bis-Tris gels (Invitrogen) and proteins were fractionated for 1-3 hrs at 64 mA in 1X running buffer (Invitrogen). PVDF membranes (Invitrogen) were pre-incubated in 100% MeOH (Thermo Fisher Scientific) for 5 min, then soaked in transfer buffer (Invitrogen) for at least 1 hour before use. Fractionated proteins in the gels were transferred electrophoretically (50 V) to the PVDF membranes overnight in a cooled chamber placed in a $4^0$C cold room.

Total transferred protein in the lysate-load was estimated via a Ponceau-S (RPI, Mt. Prospect, IL, US) staining of the membranes after transfer. Following transfer and Ponceau-S staining, the membranes were blocked in 5% milk in TBS-T, cut according to expected molecular mass, and immunoblotted overnight at 4°C with one of the following primary antibodies diluted in blocking solution: mouse monoclonal $IgG_1$ anti-HA epitope tag (26183, Invitrogen; 1:10,000-1:20,000, or 12Ca5, NeuroMab, Davis, CA, USA; 1:1000), mouse monoclonal $IgG_1$ anti-FLAG epitope tag (F1804, Sigma-Aldrich; 1:1000), or mouse monoclonal $IgG_{2a}$ anti-GFP (86/8, NeuroMab; 1:1000). Blots were rinsed twice in blocking buffer, then washed in blocking buffer three time for at least 30 min total, then incubated for 1 hr at room temp with the following secondary antibodies diluted in blocking solution: Goat anti-mouse IgG heavy chain specific (HC)-HRP conjugate (Jackson Laboratory; 1:20,000-1:40,000), goat anti-mouse IgG light chain specific (LC)-HRP conjugate (Jackson Laboratory; 1:10,000), or goat anti-mouse IgG (H+L)-HRP Conjugate (Bio-Rad, Hercules, CA, USA; 1:10,000). Following secondary incubation, blots were washed in TBS-T at least six times for at least 1 hr total, then washed in TBS three times for 10 min. The membranes were developed using Prometheus ProSignal Femto (Genesee Scientific, El Cajon, CA, USA) on autoradiography film (Amersham Hyperfilm ECL, Cytivia, Global Life Sciences Solutions USA, LLC., Marlborough, MA, USA) using a film developer.

**Cardiomyocyte staining**

Coverslips were prepared as before, then treated with poly-$_{L}$-lysine (0.01%) for 20 minutes followed by laminin (20 µg/ml) for at least 15 minutes prior to use. Freshly isolated ventricular myocytes were resuspended in a small amount of PBS, then plated on prepared coverslips and allowed to settle in a 37°C incubator for 45 minutes. For Airyscan and SMLM experiments, cultured or freshly isolated cells were treated for 8 min at 37°C with PBS (control), 100 nM ISO (Sigma-Aldrich) in PBS (ISO), 50 µM BV02 (Sigma-Aldrich) in PBS (BV02), or both 100 nM ISO and 50 µM BV02 in PBS (ISO+BV02). Cells were fixed and permeabilized with ice cold methanol (Thermo Fisher Scientific) for 5 min at -20°C, then rinsed three times with room temperature PBS, followed by three 10 min washes in PBS. Fixed cells were blocked for 1 hr at room temperature in blocking solution made with 20% fish serum blocking buffer (ThermoFisher Scientific) and 0.5% v/v Triton X-100 (Sigma-Aldrich) in PBS.

For the 14-3-3 staining comparison, plated cardiomyocyte coverslips were fixed in PFA/PEM (4% PFA, 80 mM PIPES [ThermoFisher Scientific], 5 mM EGTA, and 2 mM $MgCl_2$) followed by 100 mM glycine for 15 min at room temp, then rinsed in PBS twice for 3 min. Permeabilization with Triton-X was performed at room temperature with 0.5% Triton X-100 in PBS for 20 min, then washed three times for 10 min each in PBS. The cells were then blocked for 1 hr at room temp in 20% fish serum blocking buffer in PBS supplemented with either 0.5% Triton X-100 or 0.2% saponin (Sigma-Aldrich).

Cells for all groups were incubated overnight in the following primary antibodies diluted in the appropriate blocking solution: rabbit polyclonal IgG anti-$Ca_V1.2$ (CACNA1C, ACC-003, Alomone Labs; 0.3:100), and mouse monoclonal IgG2b anti-14-3-3 (pan) (H-8, sc-1657, Santa Cruz Biotechnology; 1:200). Covers were rinsed three times in 4% m/v skim milk in PBS, then washed three times in 4% m/v skim milk for ten minutes each, then incubated for 1 hr at room temperature in the following secondary antibodies diluted in blocking solution: Alexa Fluor 647-conjugated goat anti-mouse IgG2b (Invitrogen), CF 568-conjugated goat anti-rabbit (Sigma-Aldrich or Biotium, Fremont, CA, USA; 1:250).

**Confocal imaging**

For Airyscan super-resolution images, stained cardiomyocyte coverslips from all permeabilization conditions were mounted on coverslips using DAPI Fluoromount-G (SouthernBiotech, Birmingham, AL, USA). Cells were imaged on a Zeiss LSM 880 super-resolution microscope equipped with an Airyscan detector and a Plan-Apochromat 63x/1.40 oil DIC M27 objective. Images were acquired using Zen software. ImageJ/FIJI was used to perform a 20 pixel background substitution, threshold each channel individually, then measure the integrated density of each channel. Channels were multiplied to determine the overlapping pixels. The integrated density of the multiplied image was divided by one of the original binary images to assess the percent of that channel colocalized with the other.

To perform $Ca_V1.2$ surface expression imaging experiments, transfected tsA-201 coverslips were fixed with 4% paraformaldehyde (PFA; Electron Microscopy Sciences, Hatfield, PA, USA) in PBS for 20 min at room temperature followed by 100 mM glycine (Sigma-Aldrich) then rinsed 2 times for 3 min in PBS. Cells were imaged on a W1-spinning Disk confocal (Andor, Belfast, UK) with a Borealis modification and TIRF module, built around an IX83 inverted microscope (Olympus, Tokyo, Japan) equipped with a 60×/1.49 NA TIRF objective lens. ImageJ/FIJI was used to measure membrane and cytosol intensity.

**Proximity Ligation Assays (PLA)**

Treated cardiomyocyte coverslips were fixed with 4% paraformaldehyde in PBS for 20 min at room temperature followed by 100 mM glycine then rinsed in PBS 2 times for 3 min. Permeabilization was performed at room temperature with 0.1% Triton X-100 in PBS for 20 min, then washed three times for 10 min each in PBS. Cells were blocked for 1 hr at room temp in 20% fish serum blocking buffer containing 0.25% v/v Triton X-100 in PBS, then incubated with primary antibody overnight at 4°C. Rabbit anti-$Ca_V1.2$ (N263/31, AB_11000167, NeuroMab; 1:100) and mouse monoclonal IgG2b anti-14-3-3 (pan) (H-8, sc-1657, Santa Cruz Biotechnology; 1:200) were used to probe for 14-3-3 and $Ca_V1.2$ proximity sites. Samples were incubated for 1 hr at 37°C with PLA probes (anti-mouse MINUS and anti-rabbit PLUS; Sigma-Aldrich), ligated for 30 min, amplified for 100 min, and finally washed in Duolink buffer B, according to the manufacturer's directions (Duolink PLA kit, Sigma-Aldrich). Coverslips were mounted using Duolink *In Situ* mounting medium with DAPI (Sigma-Aldrich) and imaged on a Zeiss LSM 880 super-resolution microscope equipped with an Airyscan detector and a Plan-Apochromat 63x/1.40 oil DIC M27 objective. Images were acquired using Zen software and maximum intensity z-projections from 0.5 µm slice intervals were generated and used to measure area and quantify puncta per area in ImageJ/FIJI.

**Single Molecule Localization Microscopy (SMLM)**

Methanol-fixed stained cardiomyocyte coverslips were washed in PBS, then mounted onto glass depression slides (neoLab, Heidelberg, Germany) with an imaging buffer containing 10 parts cysteamine hydrochloride (MEA) buffer (100 mM MEA [Sigma-Aldrich] in PBS; pH 8.0 with NaOH), 89 parts buffer B (50 mM Tris pH 8.0, 10 mM NaCl, and 10% w/v glucose [Sigma-Aldrich] in water) and 1 part glucose oxidase (GLOX) oxygen scavenging system (10 mM Tris pH 8.0, 56 mg/ml glucose oxidase [Sigma-Aldrich], and 34 mg/ml catalase [Sigma-Aldrich] in PBS). Twinsil silicone-glue (Picodent, Wipperfürth, Germany) and aluminum tape (T204-1.0 – AT205; Thorlabs Inc., Newton, NJ, USA) were used to exclude oxygen from the coverslips and the completed imaging buffer was used for a maximum of 3 hrs. Cells were imaged on a Leica DMi8 microscope (Leica Microsystems, Wetzlar, Germany) in TIRF mode with a penetration depth of 150 nm, using a HC PL APO 160x 1.43 oil CORR GSD objective (Leica Microsystems). Ground state depletion was performed, and dye-blinking was elicited using a 638 nm or 561 nm/150 mW laser. Photon emission was detected with a Hamamatsu Flash 4.0 camera. 35,000 - 50,000 frames of blinking were collected with an exposure time of 10 ms using LAS X Life Science Software (Leica Microsystems). 10 nm localization maps were produced and used to measure $Ca_V1.2$ and 14-3-3 cluster areas and mean gray value, colocalization, and overlap in ImageJ/Fiji. Imaris image analysis software was used to generate cluster area vs. distance plots and representative images.

**Whole cell patch clamp**

Freshly isolated ventricular cardiomyocytes and transfected tsA-201 cells were used to record whole cell calcium currents ($I_{Ca}$) using voltage clamp. Borosilicate glass pipettes (Sutter instruments, Novato, CA, USA) fire polished to 1-3 MΩ resistance for cardiomyocytes and 3-10 MΩ resistance for tsA-201 cell were filled with internal solution (87 mM L-aspartic acid [Sigma-Aldrich], 20 mM CsCl [Thermo Fisher Scientific], 1mM $MgCl_2$, 10 mM HEPES, 10 mM EGTA and 5mM MgATP [Sigma-Aldrich], pH 7.2 with CsOH). MgATP was added fresh on the day of the experiment. Cardiomycytes were initially perfused in Tyrode's external solution (140 mM NaCl, 5 mM KCl, 10 mM HEPES, 10 mM dextrose [Thermo Fisher Scientific], 1 mM $MgCl_2$, and 2 mM $CaCl_2$, pH 7.4 with NaOH). tsA-201 cells were initially perfused in a 2 mM calcium external solution (5 mM CsCl, 10 mM HEPES, 10 mM dextrose, 140 mM NMDG, 1 mM $MgCl_2$, and 2 mM $CaCl_2$, pH 7.3 with HCl). After obtaining whole cell configuration, cardiomyocyte perfusion was switched to the same 2 mM calcium extern al solution, which was maintained through recording. For tsA-201 cells, perfusion was switched to a 20 mM calcium solution (5 mM CsCl, 10 mM HEPES, 10 mM dextrose, 113 mM NMDG, 1 mM $MgCl_2$, and 20 mM $CaCl_2$, pH 7.3 with HCl). tsA-201 baseline currents were recorded after 1-3 min to allow the solution to completely change. To prevent $I_{Ca}$ rundown in the cardiomyocytes during the treatment period, cells were held at -80 mV for 8 min prior to recording of the control current. The recording protocol was held at -80 mV, stepped to -40 mV for 100 ms to inactivate sodium channels, then stepped to voltages between -60 to 80 mV for 300 ms. Following control recordings, cells were treated with 100 nM ISO in the 2 mM calcium external solution for 3 min, and recorded again. Currents were recorded at 10kHz, filtered using a low-pass 2 kHz filter using an Axopatch 200B amplifier (Molecular Devices, Sunnyvale, CA, USA), and current-voltage relationships were plotted and fit with the Boltzmann sigmoidal equation using Prism (GraphPad Software Inc., La Jolla, CA, USA). Membrane potentials were corrected for a liquid junction potential of -10 mV. All experiments were performed at room temp. Membrane potential values in results and datasheet are corrected for the liquid junction potential.

**Single channel patch clamp**

Transfected tsA-201 cells were used to record single channel calcium currents ($I_{Ca}$) using voltage clamp. Borosilicate glass pipettes (Sutter instruments, Novato, CA, USA) were pulled to 3-10 MΩ resistance and filled with internal solution (20 mM $CaCl_2$, and 10 mM HEPES, pH to 7.2 with CsOH) supplemented with 500 nM BayK 8644. Cells were initially perfused in 2 mM calcium external solution. Upon obtaining a giga seal, perfusion was swapped to a high potassium solution (145 mM KCl, 2 mM $MgCl_2$, 0.1 mM $CaCl_2$, 10 mM HEPES, and 10 mM dextrose, pH to 7.3 with KOH). After 1 min to ensure bath solution had completely changed, single channel currents were recorded. Cells were held at 80 mV for 100 ms, then stepped to 30 mV for 2000 ms, followed by another 80 mV step for 100 ms. This protocol was repeated 50 times per cell. Currents were recorded at 20 kHz, filtered using a low-pass 2 kHz filter using an Axopatch 200B amplifier (Molecular Devices), digitized using a Digidata 1550B plus Humsilencer (Molecular Devices) and acquired using pClamp (Molecular Devices). Clampfit software (Molecular Devices) was used to detect single channel openings, measure their amplitudes, and to determine channel $P_o$.

Membrane potential values in results and datasheet are corrected for the liquid junction potential. Cooperativity was assessed with a binary coupled Markov chain model using a program run in Matlab (Mathworks, Natick, MA, USA), as previously described (4-6). $Ca_V1.2$ single channel current amplitude was set to 0.5 pA.

**Cardiomyocyte culture**

Coverslips (#1.5; VWR, Radnor, PA, USA) were sonicated 20 min in 1 M NaOH (Thermo Fisher Scientific), washed in DI water for an hour, stored in 70% ethanol, then treated with poly-$_L$-lysine (0.01%; Sigma-Aldrich) for 20 minutes followed by laminin (20 µg/ml; Life Technologies, Carlsbad, CA, USA) for at least 15 minutes prior to use. Freshly isolated cells were resuspended and plated in sterile filtered minimum essential medium (MEM; Gibco [Thermo Fisher Scientific, Waltham, MA, USA]) supplemented with 5% fetal bovine serum (FBS; Gibco), 0.1% penicillin/streptomycin (Gibco), 1% insulin transferrin selenium (ITS; Gibco), 2 mM glutamax (Gibco), 10 mM HEPES, 4 mM $NaHCO_3$ (Sigma-Aldrich), and 0.2% BSA. Cells were left in a cell culture incubator (37°C, 95:5% $O_2$:$CO_2$) for at least 1 hour, then the media was replaced with sterile filtered MEM supplemented with 0.1% penicillin/streptomycin, 1% ITS, 2 mM glutamax, 10 mM HEPES, 4 mM $NaHCO_3$, and 0.2% BSA with Ad-14-3-3ε-mRuby, Ad-EYFP-difopein, or Ad-RFP (repackaged from plasmids into adenovirus by the UC Davis CVRI Viral Vector Core) at 5 x $10^7$ virus particles per million cells. After 24 hours, the media was replaced with fresh media without adenovirus for another 24 hours before use.

**AlphaFold3 predictions of protein-protein interactions**

We used the AlphaFold 3 server to model the full sequences of human $Ca_v1.2$ $\alpha_{1C}$ subunit (Uniprot ID: Q13936), human $Ca_v\beta_2$ subunit (Uniprot ID: Q08289), rabbit $Ca_v1.2$ $\alpha_{1C}$ subunit (Uniprot ID: P15381), rabbit $Ca_v\beta_2$ subunit (Uniprot ID: P54288), incorporating post-translational modifications at S1700 and S1928 for rabbit $Ca_v1.2$, and S1718 and S1981 for human $Ca_v1.2$, and human 14-3-3ε (Uniprot ID: P62258). For each model, we generated 100 predictions and ranked them based on pLDDT scores of the full-length $Ca_v1.2$ α subunit.

**Statistical analysis**

*N* represents the number of animals and *n* represents the number of cells. Data are reported as mean ± SEM. Statistics were performed using Prism (GraphPad Software Inc.), and all data sets were tested for normality. Unpaired Student's t-tests or Mann Whitney tests were used to compare data sets with two groups, one-way ANOVAs or Krushkal-Wallis (non-parametric) tests with post-hoc testing were used to compare data sets with more than two groups, or two-way ANOVAS when there were two independent variables. $P < 0.05$ was considered statistically significant.

### Supplemental Figures and Legends

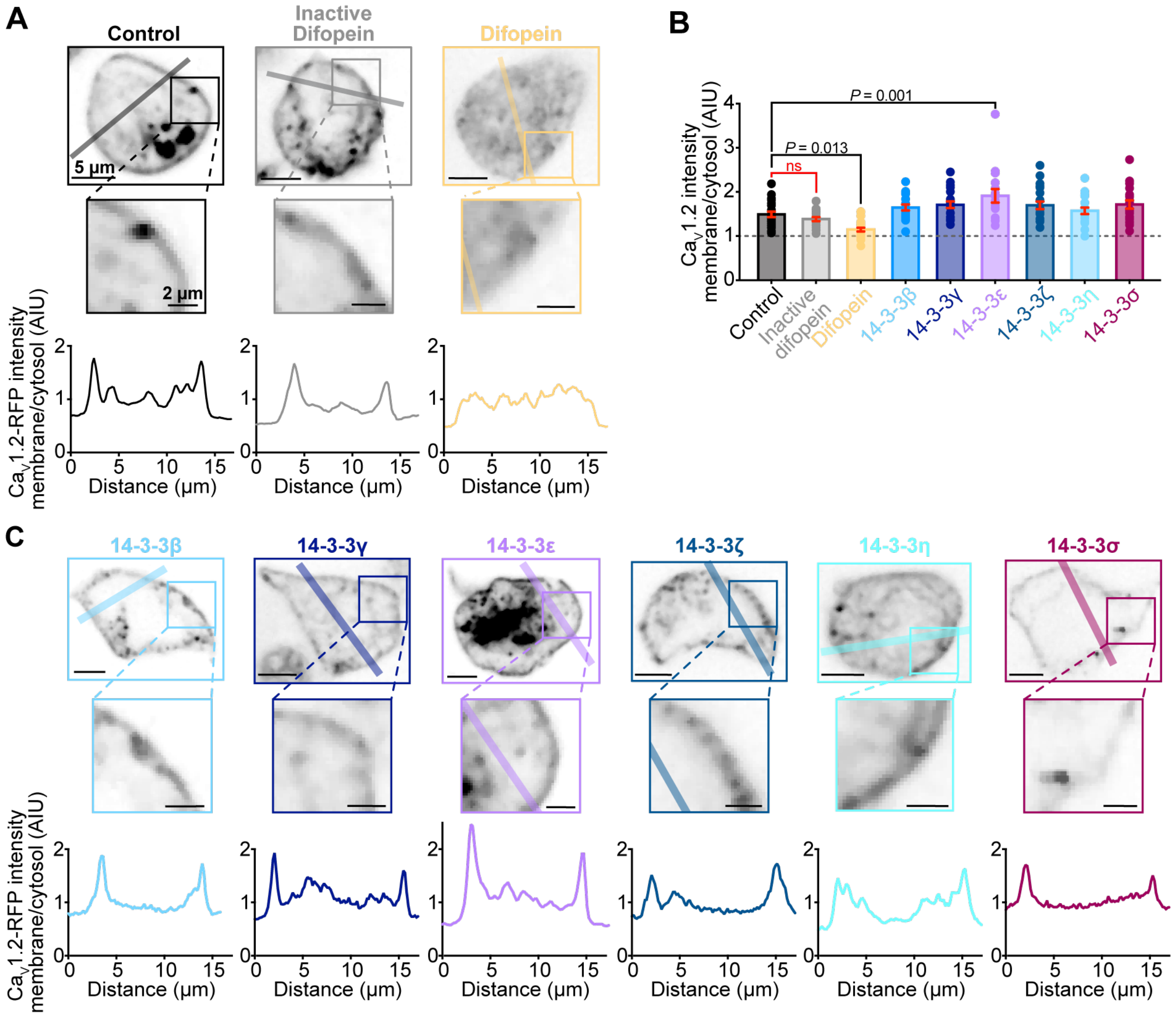


**Figure S1. Ca$_V$1.2 surface expression is altered by 14-3-3ε overexpression or competitive inhibition in tsA-201 cells. A**, representative confocal images of tsA-201 cells transiently transfected with GFP-Ca$_V$1.2 and auxiliary subunits alone (*control*), or co-transfected with inactive 14-3-3 peptide competitive inhibitor difopein (*inactive difopein*), active difopein (difopein). **B**, histogram summarizing the ratio between membrane and cytosolic GFP intensity (difopein: $n$ = 20; inactive difopein: $n$ = 18; control: $n$ = 22; 14-3-3η: $n$ = 20; 14-3-3β: $n$ = 19; 14-3-3ζ: $n$ = 21; 14-3-3γ: $n$ = 20; 14-3-3σ: $n$ = 19; 14-3-3ε: $n$ = 15). **C,** representative confocal images of tsA-201 cells transiently transfected with GFP-Ca$_V$1.2 and auxiliary subunits and overexpressing the indicated 14-3-3 isoform. Colored bars indicate regions used for the below line scan showing the ratio between membrane and cytosolic GFP intensity in each case. Data were analyzed using one-way ANOVAs with multiple comparison post-hoc tests. Data are presented as mean ± SEM. Scale bars indicate 5 µm for full images and 2 µm for zoomed in regions.

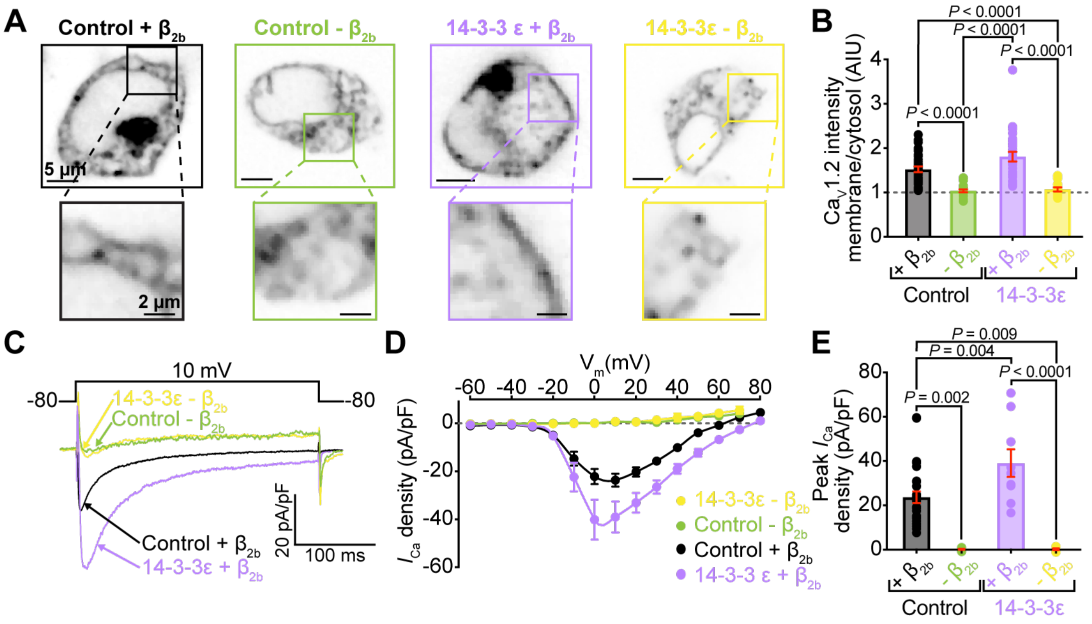


**Figure S2. Overexpression of 14-3-3ε is not sufficient to drive trafficking of $Ca_V1.2$ to the membrane in the absence of $Ca_V\beta_{2b}$ auxiliary subunits. A**, representative confocal images of tsA-201 cells transiently transfected with RFP-$Ca_V1.2$ and $Ca_V\alpha_2\delta$ auxillary subunit alone (*-$\beta_{2b}$ control*), or co-transfected with $Ca_V\beta_{2b}$ (*+$\beta_{2b}$ control*), with overexpressed 14-3-3ε (*-$\beta_{2b}$ 14-3-3ε*), or both $Ca_V\beta_{2b}$ and 14-3-3 (*+$\beta_{2b}$ 14-3-3ε*). Scale bars indicate 5 µm for full images and 2 µm for zoomed in regions. **B**, histogram summarizing the ratio between membrane and cytosolic GFP intensity (-$\beta_{2b}$ control: *n* = 20; +$\beta_{2b}$ control: *n* = 41; -$\beta_{2b}$ 14-3-3ε: *n* = 18; *+$\beta_{2b}$ 14-3-3ε*: *n* = 34). **C,** representative whole-cell currents elicited from transfected tsA-201 cells (-$\beta_{2b}$ control; lime, +$\beta_{2b}$ control; black, -$\beta_{2b}$ 14-3-3ε; yellow, *+$\beta_{2b}$ 14-3-3ε*; lavender). **D,** plots showing the voltage dependence of $I_{Ca}$ density for all groups (-$\beta_{2b}$ control: *n* = 7; +$\beta_{2b}$ control: *n* = 14; -$\beta_{2b}$ 14-3-3ε: *n* = 5; *+$\beta_{2b}$ 14-3-3ε*: *n* = 9). **E**, histogram showing peak $I_{Ca}$ density for all groups shown in D. Data were analyzed using two-way ANOVAs with multiple comparison post-hoc tests. Data are presented as mean ± SEM.

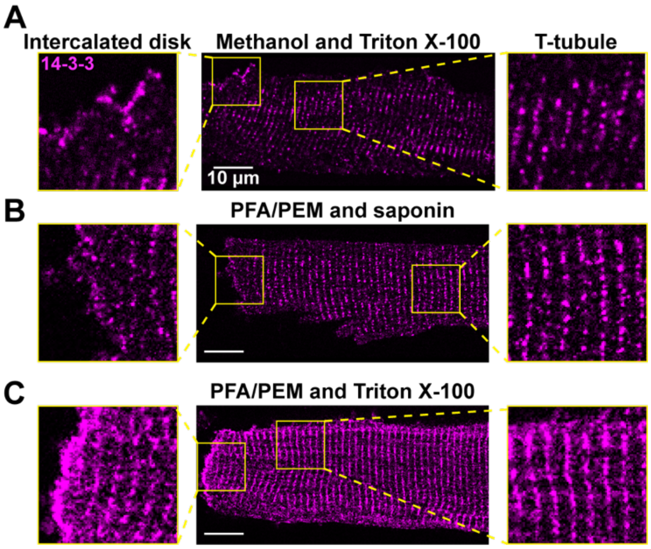


**Figure S3. 14-3-3 localization is preserved across permeabilization conditions.** Representative Airyscan images of immunostained 14-3-3 in myocytes fixed with **A**, methanol and permeabilized with Triton X-100; **B**, PFA/PEM and permeabilized with Triton X-100; or **C**, PFA/PEM and permeabilized with saponin. Scale bars indicate 10 µm.

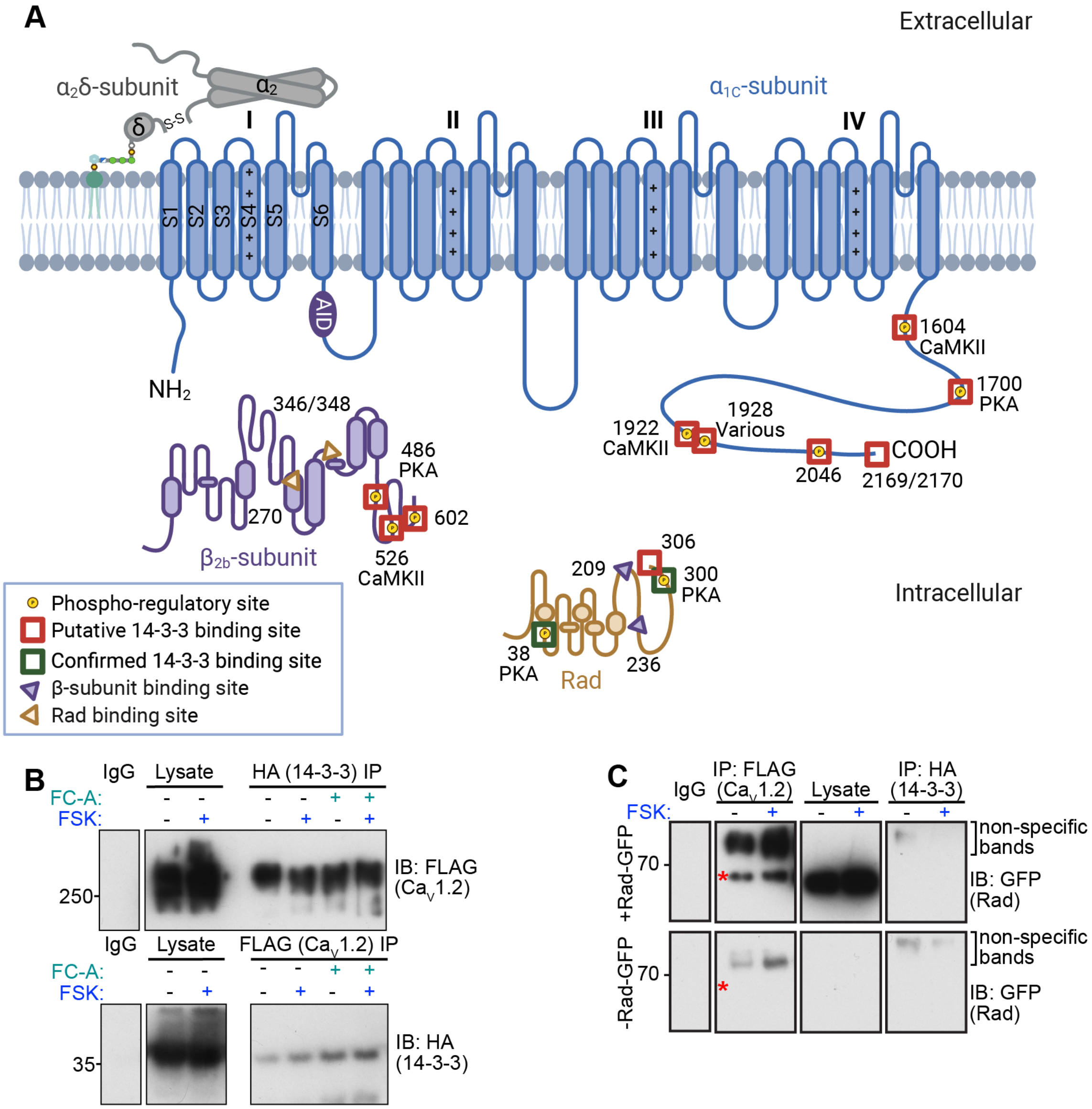


**Figure S4. Potential 14-3-3 binding sites. A,** Graphical depiction of Ca$_V$α$_{1C}$ (UniProtID: P15381-1), Ca$_V$β$_2$ (P54288-1), and Rad (P55042) structures showing known or suspected phosphorylation sites (yellow dots), overlayed with potential 14-3-3 binding sites (red boxes), or known 14-3-3 binding sites (green boxes). Created with BioRender.com. **B,** western blot showing lysate, 14-3-3 IP, and Ca$_V$1.2 IP lanes probed for Ca$_V$1.2 (top), or probed for 14-3-3 (bottom) with and without forskolin treatment (FSK, blue), and treatment with 14-3-3 mode III stabilizer fusiccocin-A (FC-A, teal). **C,** western blot showing lysate, 14-3-3 IP, and Ca$_V$1.2 IP lanes probed for GFP (Rad) in cells transfected with Rad (top), or cells not transfected with Rad (bottom) with and without forskolin treatment (FSK, blue). Red asterisks indicate expected molecular weight of Rad-GPF. See Supplemental Information for uncropped scans.

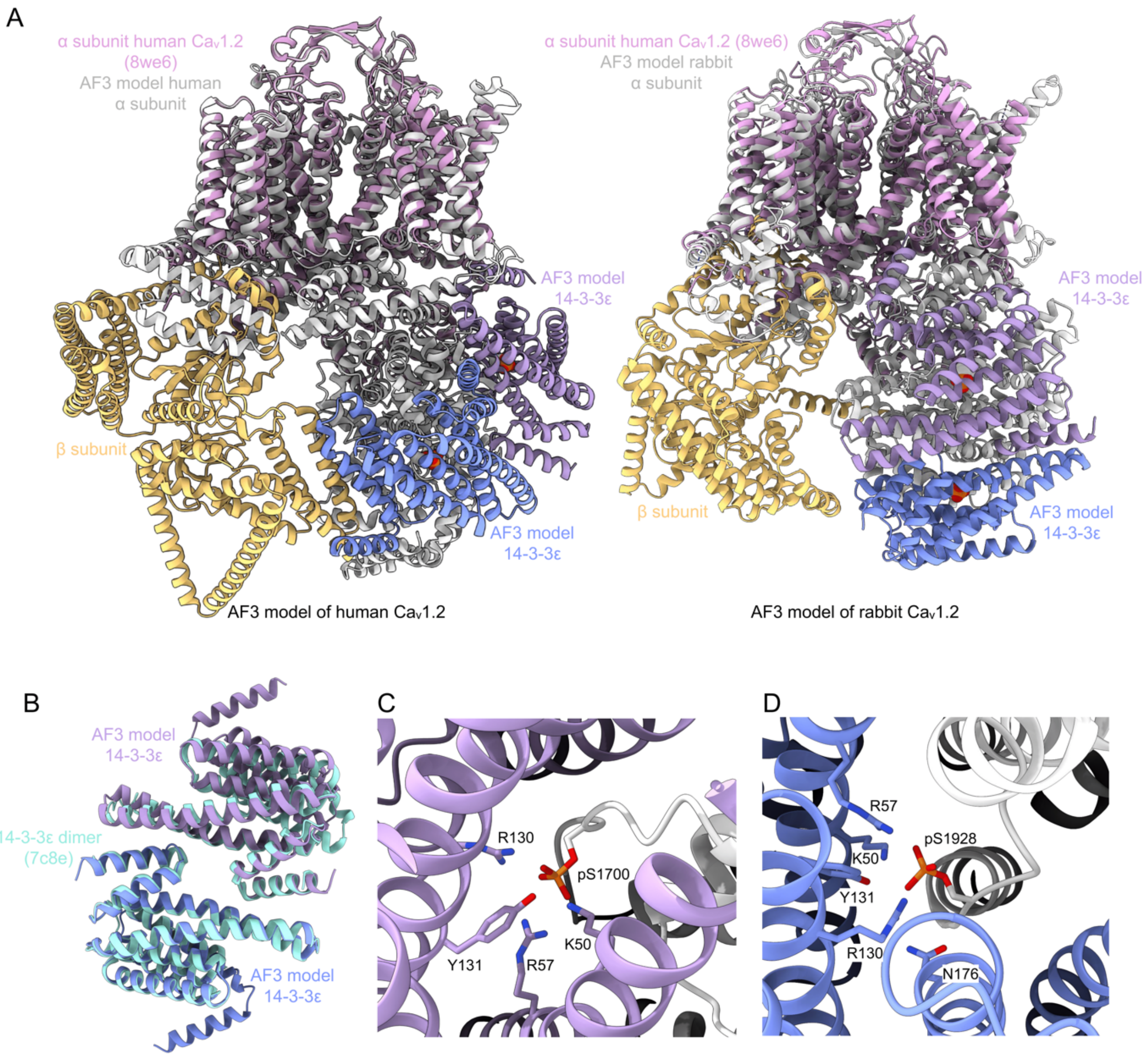


**Figure S5. AF3 models of $Ca_V$1.2 and 14-3-3ε dimer. A**, AF3 model of human (left panel) and rabbit (right panel) $Ca_V$1.2 $α_{1C}$ (colored in light gray), $Ca_V$1.2 $β_{2b}$ (colored in yellow) subunits, and 14-3-3ε dimer (individual subunits colored in blue and purple), superimposed with structure of human $Ca_V$1.2 $α_{1C}$ (PDB ID: 8we6, in pink) with an RMSD of ~2.8 Å and ~3.2 Å, respectively. Phosphorylation sites are shown in sphere representation. **B**, AF3 model of human 14-3-3ε dimer (colored in blue and purple) superposed with x-ray structure of 14-3-3ε (PDB ID: 7c8e, colored in cyan) with a RMSD of ~0.9 Å. **C**, Close-up view of pS1700 interactions with 14-3-3ε residues. Side chains of pS1700 and key 14-3-3ε residues are shown in stick representation. **D**, Close-up view of pS1928 interactions with 14-3-3ε residues. Side chains of pS1928 and key 14-3-3ε residues are shown in stick representation.

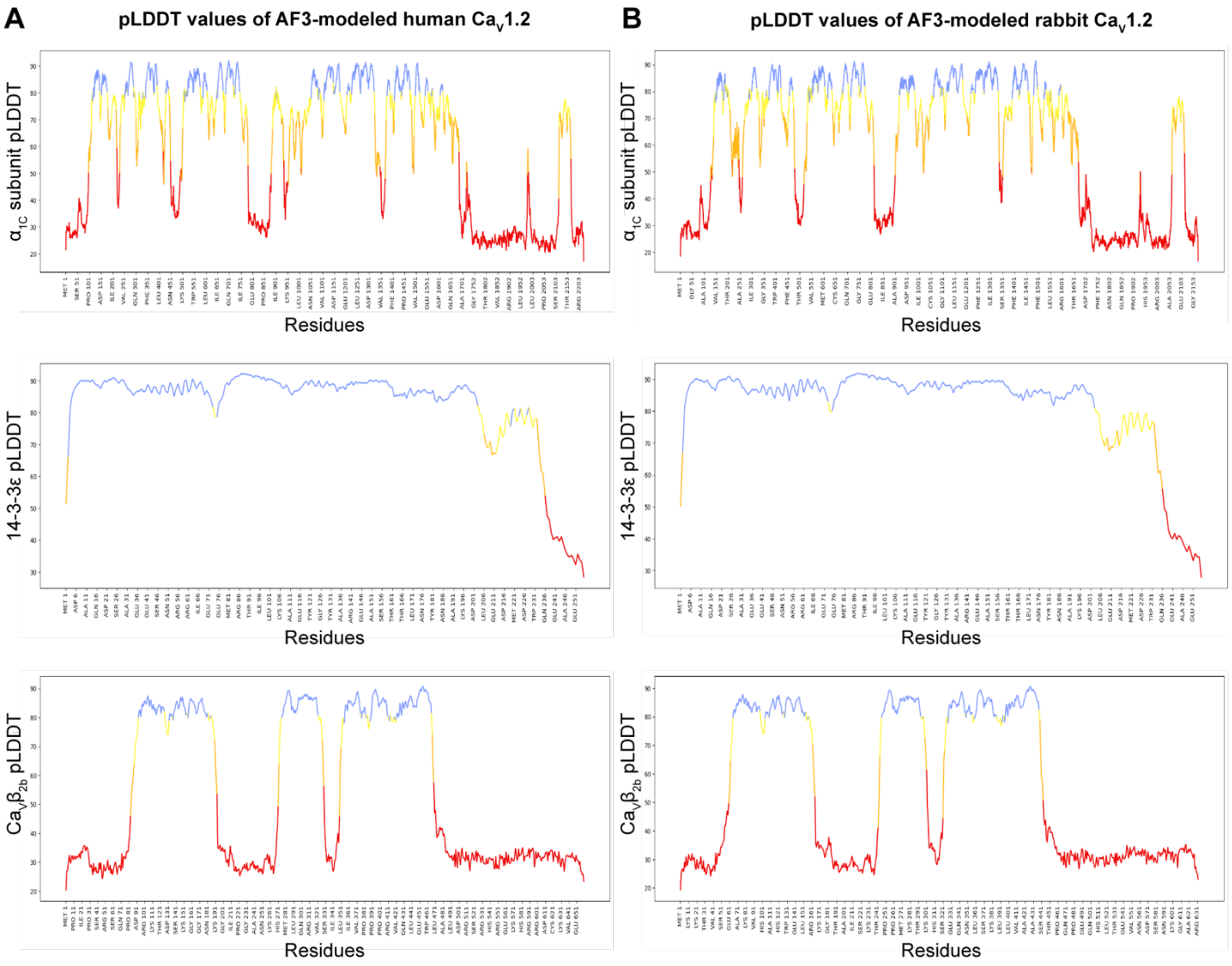


**Figure S6. Per-residue predicted Local Distance Difference Test (pLDDT) values of AF3 models. A**, pLDDT values for AF3 models of human Ca$_V$1.2 $\alpha_{1C}$ (top panel), 14-3-3ε (middle panel), and Ca$_V$1.2 $\beta_{2b}$ (bottom panel). **B**, pLDDT values for AF3 models of rabbit Ca$_V$1.2 $\alpha_{1C}$ (top panel), 14-3-3ε (middle panel), and Ca$_V$1.2 $\beta_{2b}$ (bottom panel).

**Supplemental Tables and Legends**

| | $V_{1/2}$ | $V_{1/2}$ comparisons | Slope | Slope comparisons |
|---|---|---|---|---|
| **Control** | -2.10 ± 2.57 | Control vs Difopein $^{ns}$ | 6.92 ± 0.50 | Control vs Difopein $^{P < 0.0001}$ |
| **14-3-3ε** | -5.25 ± 2.95 | 14-3-3ε vs control $^{ns}$ | 4.54 ± 0.76 | 14-3-3ε vs control $^{P = 0.045}$ |
| **Difopein** | 4.37 ± 4.13 | Difopein vs 14-3-3ε $^{ns}$ | 11.96 ± 0.88 | Difopein vs 14-3-3ε $^{P < 0.0001}$ |

**Table S1. tsA-201 cell voltage dependence of $G/G_{max}$.** Mean ± SEM values for voltage at half activation ($V_{1/2}$) and slope factor obtained after fitting the voltage dependence of $G/G_{max}$ in each condition with a Boltzmann function. Data were compared using one-way ANOVAs with multiple comparison post-hoc tests.

| | $V_{1/2}$ | | Slope | |
|---|---|---|---|---|
| | Control | ISO | Control | ISO |
| **RFP** | -18.08 ± 3.13 | -29.94 ± 2.39. $^{P < 0.0001}$ | 6.82 ± 0.52 | 5.15 ± 0.32 $^{P = 0.0003}$ |
| **14-3-3ε** | -19.93 ± 2.15 | -30.22 ± 1.23 $^{P < 0.0001}$ | 5.61 ± 0.28 | 4.56 ± 0.38 $^{P = 0.028}$ |
| **Difopein** | -20.41 ± 1.72 | -28.75 ± 1.59 $^{P < 0.0001}$ | 5.98 ± 0.11 | 5.78 ± 0.30 $^{ns}$ |

**Table S2. Cultured cardiomyocyte voltage dependence of $G/G_{max}$.** Mean ± SEM values for voltage at half activation ($V_{1/2}$) and slope factor obtained after fitting the voltage dependence of $G/G_{max}$ in each condition with a Boltzmann function. Data were compared using two-way ANOVAs with multiple comparison post-hoc tests. *P*-values correspond to comparisons between control and ISO in each group.
